# Structural basis for the ATP-dependence of omegasome biogenesis by DFCP1

**DOI:** 10.64898/2026.05.27.728090

**Authors:** Mohit Misra, Zhicheng Cui, Marie-Catherine Drigeard Desgarnier, Amin Godin, Deep Chatterjee, Yuchao Zhang, Julia Pomirska, Laura Rodriguez de la Ballina, Karla Tapia, Sebastian Mathea, James H. Hurley, Kay O. Schink, Ivan Dikic, Harald Stenmark, Viola Nähse

## Abstract

Autophagosome formation begins at phosphatidylinositol 3-phosphate-enriched endoplasmic reticulum (ER) subdomains termed omegasomes. DFCP1/ZFYVE1 is recruited to omegasomes through its FYVE domains and has recently been shown to function as an ATPase involved in omegasome constriction and autophagosome biogenesis. However, the structural basis of DFCP1 ATPase activity and how nucleotide-dependent conformational states regulate omegasome dynamics remain unknown. Here, we determined the crystal structures of the DFCP1 ATPase domain in complex with either ADP or the non-hydrolyzable ATP analogue, AppNHp at near atomic resolution. We employed structure-guided mutagenesis to define residues required for DFCP1 function in cells. The structures reveal that the active site contains a trans-acting Arg271 finger coordinating the γ-phosphate and trans-acting His323 that stacks on the ATP adenine ring. ATP hydrolysis results in a conformational switch in the vicinity of Arg271, while contacts with His323 persist in the presence of ADP. Biochemical analyses of DFCP1 and its mutants show that DFCP1 forms ATP-dependent microdomains on membranes. Disruption of His323 and Arg271 uncouples nucleotide binding from ATP hydrolysis and alters the oligomeric state of the protein. Live-cell imaging of DFCP1 knockout cells reconstituted with wild-type or mutant DFCP1 further demonstrates that these biochemical defects translate into distinct omegasome phenotypes. Together, our data provides a structural-function framework for DFCP1 ATPase activity and reveals how distinct catalytic elements control omegasome dynamics *in vivo*. We propose that nucleotide-dependent assembly and hydrolysis-driven conformational changes enable DFCP1 to regulate omegasome formation and its progression toward autophagosome closure.

**Significance Statement:** DFCP1 organizes the ER subdomains known as omegasomes, which are sites of autophagosome biogenesis. DFCP1 is an ATPase whose catalytic activity is required for function, but the precise role of its ATPase activity is unknown. Crystal structures of the DFCP1 ATPase domain in the presence of a non-hydrolyzable ATP analogue and ADP reveal an Arg finger that operates in trans, such that dimerization is required for ATP hydrolysis, and ATP hydrolysis destabilizes dimers. Dimerization is also promoted by an adenine base-stacking interaction in trans with H323. These residues are important for DFCP1 clustering on membranes and omegasome constriction, clarifying that the role of ATP is to promote the dimerization of the ATPase domain and thereby control the organization of DFCP1 on membranes.

## Introduction

Macroautophagy, hereafter referred to as autophagy, is a conserved lysosomal degradation pathway that removes cytoplasmic material, damaged organelles, protein aggregates, and invading pathogens. During autophagy, cargo is engulfed by a newly formed double-membrane phagophore, which subsequently fuses with lysosomes for degradation (Melia et al., 2020). Phagophore growth requires rapid lipid supply and occurs in close proximity to the endoplasmic reticulum (ER), the major lipid-synthesizing organelle of the cell (Maeda et al., 2019; Osawa et al., 2022). In mammalian cells, autophagosome formation starts at specialized ER-associated sites called omegasomes (Axe et al., 2008). These PI3P-enriched structures act as platforms for phagophore nucleation and expansion. Omegasomes are highly transient structures. They expand together with the phagophore, and at the maximum omegasome diameter, the phagophore extrudes out of the ring and the omegasome subsequently constricts, thereby closing the phagophore (Nahse et al., 2024).

DFCP1/ZFYVE1 is a double FYVE-domain protein that binds phosphatidylinositol 3-phosphate (PI3P) and rapidly accumulates at omegasomes upon autophagy induction. For many years, DFCP1 was mainly used as a marker of early autophagosome formation (Axe et al., 2008). However, we recently showed that DFCP1 is an active regulator of omegasome dynamics. DFCP1 functions as an ATPase, dimerizes in a nucleotide-dependent manner, and is required for omegasome constriction and autophagosome closure (Nahse et al., 2023). The structural basis of DFCP1 ATPase function, however, remained unknown.

Nucleotide-dependent membrane remodeling is a common principle in cell biology. ATPases and GTPases use cycles of nucleotide binding, hydrolysis, oligomerization, and conformational change to remodel membranes, organize membrane-associated assemblies, and drive membrane constriction or fission (Daumke and Praefcke, 2016; Jimah and Hinshaw, 2019). Although DFCP1 is distinct from canonical membrane-remodeling NTPases, its ATPase activity suggests that omegasome dynamics may also be controlled by nucleotide-dependent structural transitions.

Here, we report crystal structures of the DFCP1 ATPase domain in complex with ADP and AppNHp at 1.9 Å and 2.3 Å resolutions, respectively. The structures reveal the nucleotide-binding pocket and dimerization interface, and identify conserved residues that support ATP binding, ATP hydrolysis, and ATP-dependent dimerization. Comparison of ADP- and ATP-bound states reveals a nucleotide-dependent conformational switch. Guided by the structure, we generated DFCP1 point mutants targeting the arginine finger, and the histidine stack. We reconstituted DFCP1 knockout U2OS cells with wild-type or mutant DFCP1 and analyzed omegasome dynamics by live-cell imaging. All ATPase-domain mutants altered omegasome dynamics, but they produced distinct phenotypes, indicating that different structural elements of the ATPase domain control different steps of omegasome maturation.

Together, our findings define the structural basis of DFCP1 ATP catalysis and show how nucleotide binding, dimerization and hydrolysis regulate DFCP1 function at omegasomes. This study provides a molecular blueprint for understanding how DFCP1 ATPase activity controls omegasome maturation and autophagosome biogenesis.

## Results

### DFCP1 ATPase domain adopts distinct ADP- and ATP-bound conformations

To define the structural basis of DFCP1 ATPase activity, we crystallized the human DFCP1 ATPase domain spanning residues 146–409 in nucleotide-bound states and solved structures with ADP and the non-hydrolysable ATP analogue AppNHp. The ADP-bound crystals diffracted at a higher resolution of 1.9 Å whereas the AppNHp bound crystals diffracted to 2.3 Å resolution (Table 1). The structures revealed a dimeric ATPase domain with one nucleotide-binding pocket in each monomer (Fig.1A). Superposition of the ADP- and AppNHp-bound structures showed the RMSD of 1.76 Å, suggesting that the overall fold is similar, however, the nucleotide hydrolysis induces local conformational changes around the active site and dimer interface. The ADP bound crystals were packed in monoclinic P 1 21 1 whereas AppNHp bound crystals were found in higher symmetry orthorhombic P21 2 21 space group. ADP bound structure had four chains of DFCP1 in the asymmetric unit while AppNHp bound structure has three molecules in the asymmetric unit. The bound nucleotides were well resolved as represented by the omit maps of ADP and AppNHp (Fo-Fc electron density at 3 σ level) (Fig.1B and C). In both structures, Mg²⁺ and water molecules were positioned close to the phosphate groups and Mg²⁺ was coordinated in the typical octahedral geometry with water, phosphate or DFCP1 amino acids (Fig.1D and E).

**Figure 1.**
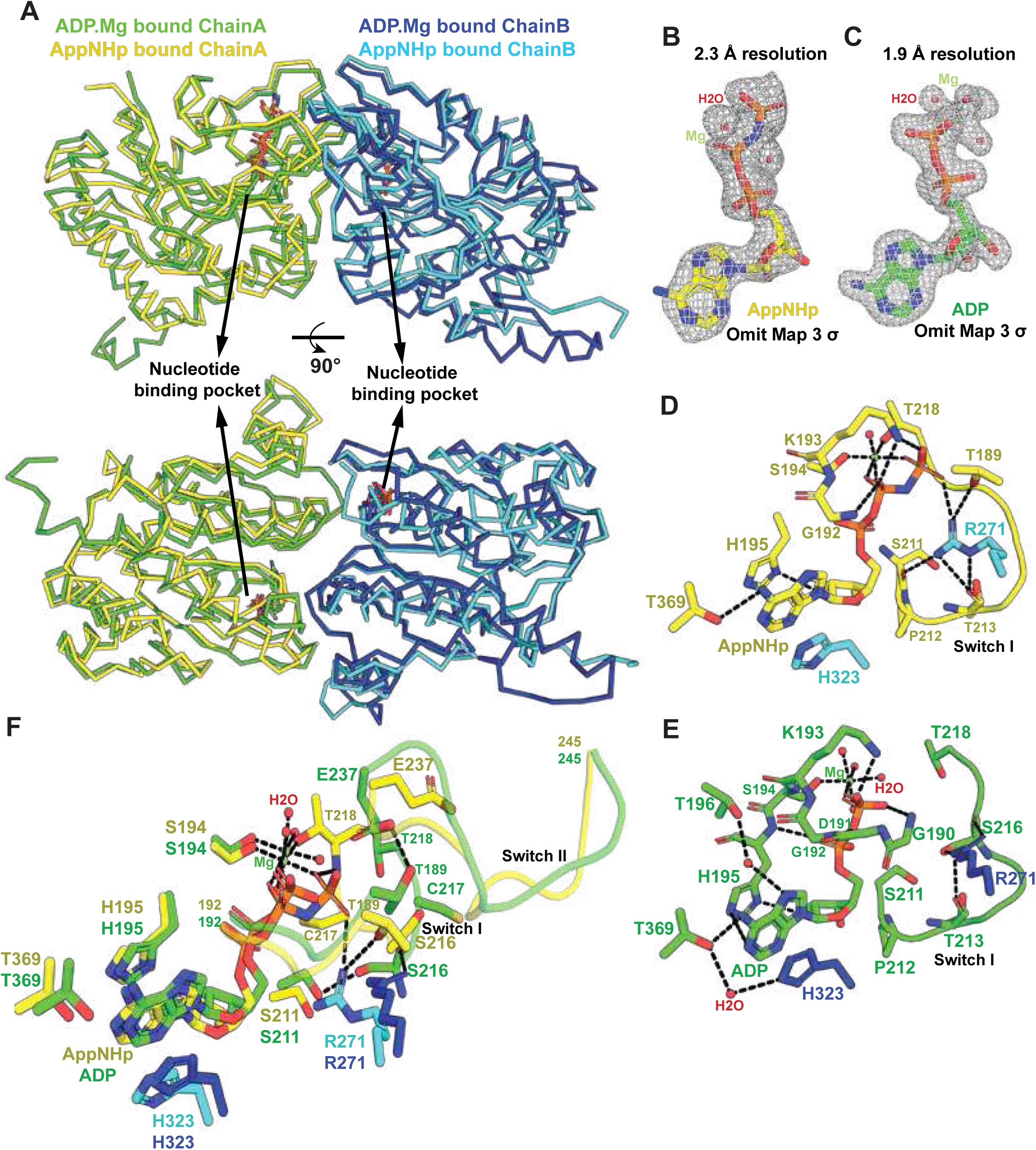
Crystal structures of DFCP1 ATPase domain bound to either ADP or AppNHp. (A) Superimposition of dimers (Chain A and Chain B) from crystal structures of DFCP1-ADP complex and AppNHP bound state. The nucleotide binding pockets are highlighted with arrows. Omit maps (Fo-Fc at 3 α) for AppNHp (B) and ADP (C) show clear occupancy of the ligands. Zoom-in view of the nucleotide binding pocket for AppNHp bound form (D) and ADP complex (E) with the key H-bonds highlighted with dashed lines. (F) Overlay of substrate (AppNHp) and product (ADP) bound states highlighting the conformational plasticity in switch I and switch II as well as key interactions with dashed lines.

**Table 1.**
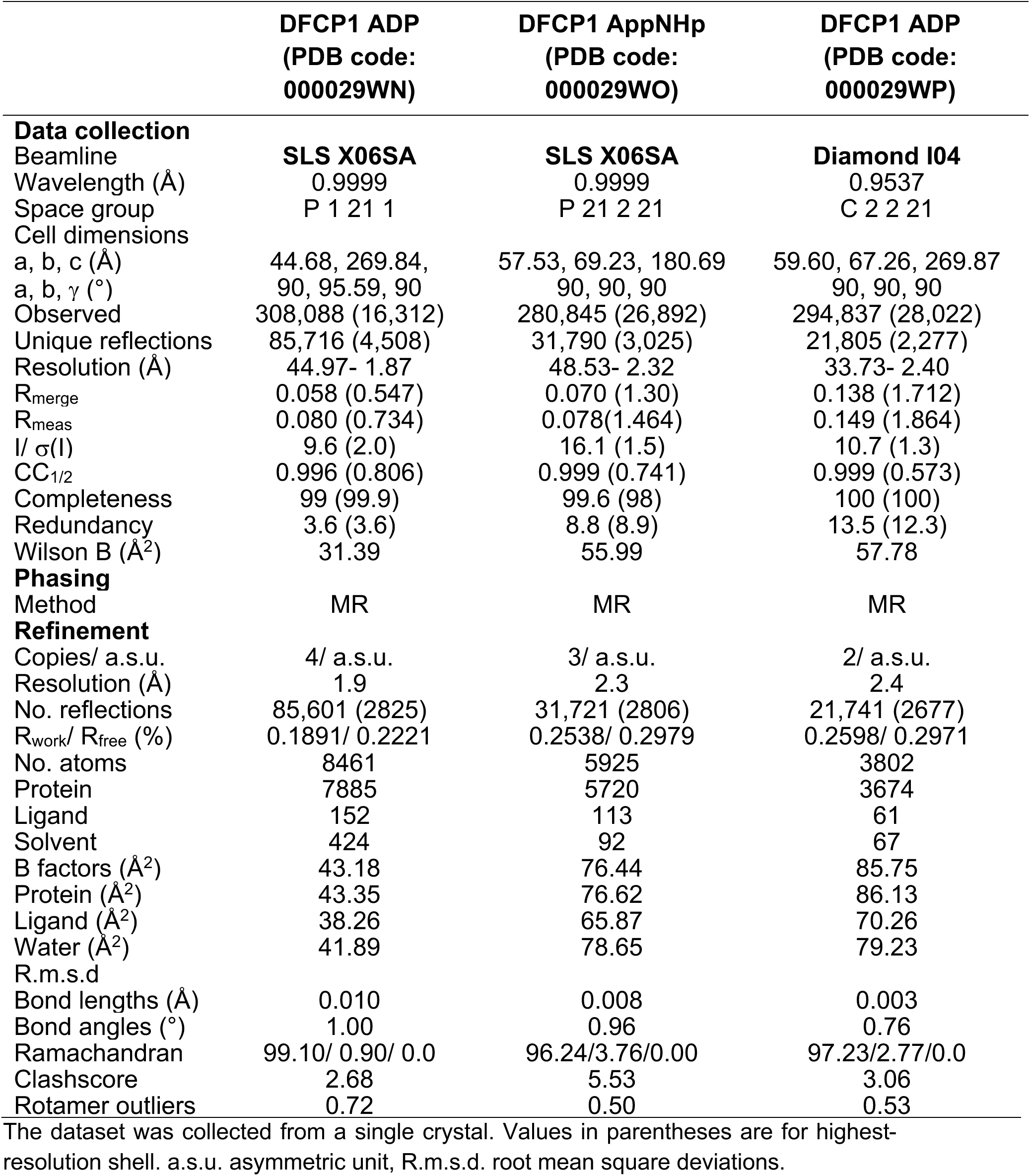
Data collection and refinement statistics for DFCP1 crystal structures.

The nucleotide-binding pocket is formed by conserved residues surrounding the phosphate groups and the ribose-base region. Close inspection of the nucleotide-binding pocket of chain A in the AppNHp-bound state revealed a strong resemblance to the phosphate-binding core of P-loop NTPases. The GDGKS segment forms a Walker A-like loop, with K193 stabilizing the terminal phosphates and S194 and T218 contributing to Mg²⁺-phosphate coordination. Unlike canonical Ras/EF-Tu GTPases, where guanine specificity and nucleotide-state switching are mediated by conserved GTPase motifs and switch regions, this site appears adapted for adenine nucleotide recognition. The adenine base is stabilized by a local polar network and a notable H195–adenine π-stacking interaction (Fig.1D).

Notably, two residues from the neighbouring protomer, chain B, also contact the AppNHp molecule bound to chain A. R271 from the opposing monomer reaches into the active site and contacts the terminal phosphate region as well as residues T189, S211 and T213. This identifies R271 as a trans-acting arginine finger, which has been observed in AAA+ATPases and other oligomeric NTPases (Hanson and Whiteheart, 2005), consistent with a role in ATP hydrolysis at the dimer interface. In addition, H323 from the opposing protomer lies close to the adenine region, together with H195 from the nucleotide-binding monomer. These two histidines sandwich the nucleotide from opposite sides of the dimer interface, forming a histidine stack that, to our knowledge, has not been described in other NTPases. This arrangement suggests that both H195 and H323 help stabilize nucleotide binding and/or promote formation of the active dimeric state.

In the ADP-bound state, the interaction between R271 and the γ-phosphate is lost, and R271 instead interacts with S216 and G190. This interaction places the switch I loop, comprising residues 214–218, in an open conformation in the absence of the γ-phosphate (Fig. 1E). Mg²⁺ coordination is also altered in the ADP-bound state: T218, which contacts Mg²⁺ in the AppNHp-bound state, flips away from the nucleotide-binding pocket, and its position is occupied by a water molecule. Owing to the higher resolution of the ADP-bound structure, we also identified several water-mediated hydrogen bonds, including interactions between T196 and the adenine base, and between H323 of the neighbouring monomer and T369.

Comparison of the AppNHp- and ADP-bound structures revealed conformational changes around switch I and switch II (Fig.1F). In the ATP-like AppNHp state, switch I is positioned close to the nucleotide and the trans-acting R271. In the ADP-bound state, the corresponding region adopts a different conformation, while R271 and T218 are repositioned relative to the nucleotide-binding pocket. In addition, T189, which forms a hydrogen bond with R271 in the AppNHp-bound state, reorients in the ADP-bound state and instead forms a hydrogen bond with E237 in the switch II loop. Together, these changes indicate that ATP hydrolysis is coupled to structural rearrangements at the dimer interface.

We solved three crystal structures of the DFCP1 ATPase domain: two ADP-bound structures and one AppNHp-bound structure. The structures crystallized in different space groups. The ADP-bound crystals belonged to the monoclinic space group P2₁ and the C-centered orthorhombic space group C222₁, whereas the AppNHp-bound crystals belonged to the primitive orthorhombic space group P2₁22₁ (Primitive orthrombic) (SI appendix, Fig. S1; Table 1). Visualization of symmetry mates in these crystal forms revealed additional intermolecular contacts beyond the ATP-dependent dimer interface. Such contacts may be relevant for understanding possible structural rearrangements during ATP hydrolysis or stabilization of higher-order oligomers. The consistency of results in difference space groups support the robust conclusions about the nature of the dimer.

Together, these structures confirm that DFCP1 dimerizes in a nucleotide-dependent manner. Both monomers contribute catalytic elements to the active site, including a trans-acting arginine finger and a unique histidine stack that position the nucleotide at the dimer interface. These features provide a structural explanation for the nucleotide-dependent dimerization and ATPase activity of DFCP1.

### Structure-guided mutagenesis impairs nucleotide-dependent DFCP1 dimerization and ATPase activity

To test how nucleotide binding affects the oligomeric state of DFCP1, we analyzed the purified DFCP1 ATPase domains by size-exclusion chromatography in the presence of ADP or the non-hydrolysable ATP analogue AppNHp. Wild-type DFCP1 shifted to an earlier elution volume (0.3 ml) in the presence of AppNHp compared with ADP, indicating that the ATP-bound state promotes formation of a larger DFCP1 species consistent with dimerization (Fig.2A). This is consistent with previous observation of nucleotide-dependent dimerization of the DFCP1 ATPase domain (Nahse et al., 2023).

**Figure 2.**
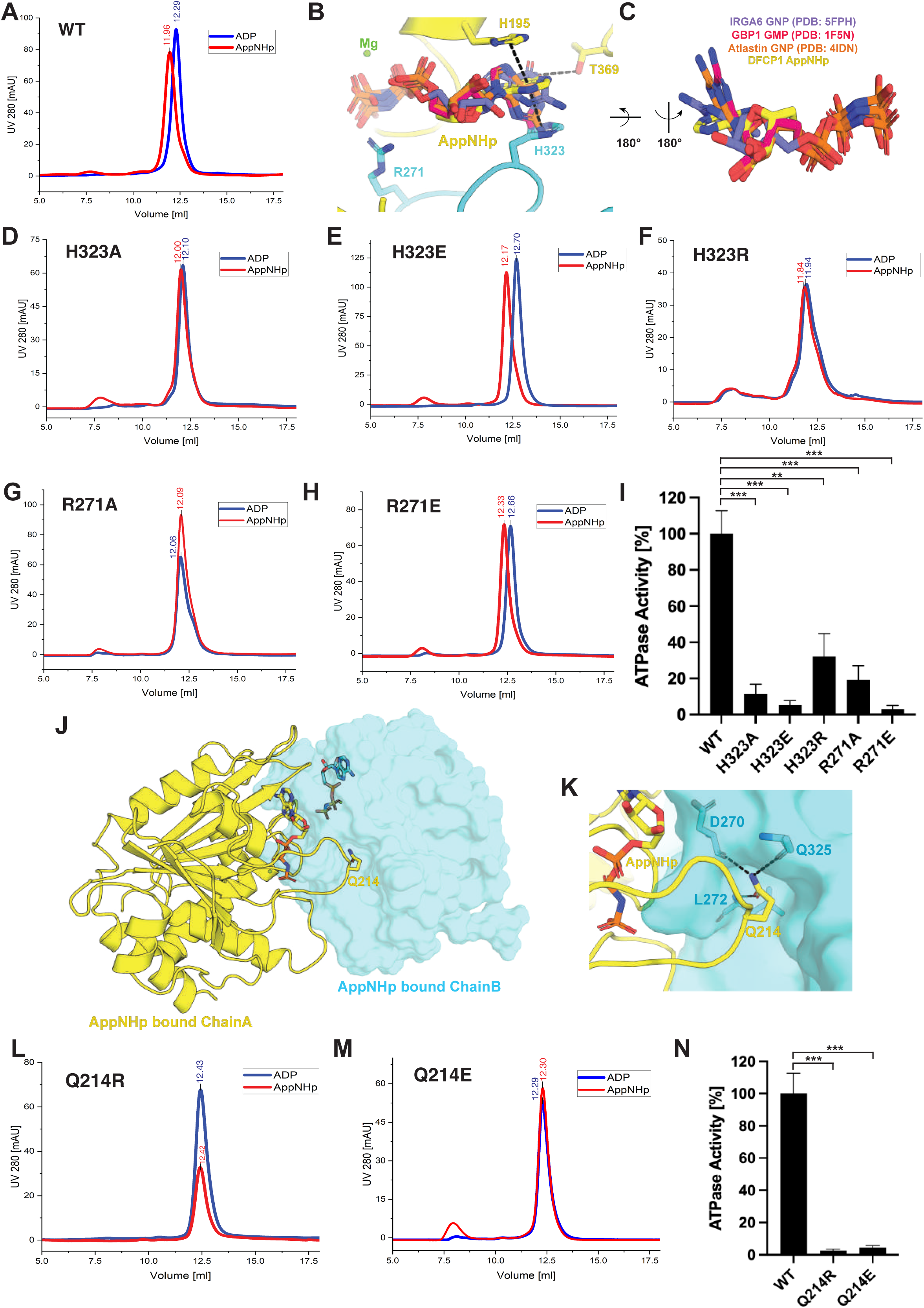
Structure-guided mutants impair nucleotide-dependent DFCP1 dimerisation and ATPase activity. (A) Size exclusion chromatography for WT DFCP1 ATPase domain shows dimer peak in the presence of AppNHp. (B) Close up view of AppNHp bound structure highlighting histidine stack and clash from guanine base superimposed from the crystal structures of Dynamin family proteins IRGA6 (PDB code: 5FPH); GBP1 (PDB code: 1F5N) and Atlastin-1 (PDB code: 4IDN). (C) Orientation of guanine base when compared to AppNHp in DFCP1 crystal structure. Results of size exclusion chromatography for mutants (D) H323A (E) H3232E (F) H323R (G) R271A (H) R271E (I) ATPase activity of mutants when compared to WT ATPase domain in ADP-Glo Kinase assay (n=3); P ≤ 0.0009 (***) and P ≤ 0.0044(**). (J) AppNHp bound Chain A and Chain B from crystal structure showing the nucleotide binding pockets relative to residue Q214 (K) Interactions of Q214 side chain with the residues in the opposing monomer. Nucleotide-dependent dimerization assay for (L) Q214R and (M) Q214E mutants. (N) ATPase activity of Q214R and Q214E mutation in comparison to WT ATPase domain(n=3); P ≤ 0.0003 (***).

Upon close examination of the adenine binding pocket, we recognized three critical interactions which support adenine binding. These include the π-stacking of H195 and H323 as well as H-bond from T369 (Fig. 2B). Dynamin family of GTPases is thought to be closely related to DFCP1 which also consists of members like GBP1, Atlastin-1 that remodel membranes with mechanical energy derived from their GTPase activity. Among its family members, GBP2 and Atlastin-3 share 23% and 24% identity with DFCP1 ATPase domain. Interestingly, H195, H323, R271 and T369 are not present in these GTPases (SI appendix, Fig. S2). Superimposition of the GNP/ GMP ligands from Dynamin family of GTPases (GBP1, Atlastin-1 and IRGA6) to the AppNHp of DFCP1 causes a serious clash from the guanine moiety of GTP to the H323 π-stacking (Fig. 2B and C). In addition, the T369 residue clashes with the oxy-group at the C6 atom of guanine while it makes favourable interaction with the amino group of adenine at similar position. The water mediated H-bond between H323 and T369 seen in the ADP bound structure further underlines the role of these residues in maintaining the adenine binding pocket (Fig. 1E). This finding suggests that that the histidine stack and T369 together act as gatekeeper residues for ATP selectivity of DFCP1. Alignment of the DFCP1 ATPase domain with closely related ATPases, including KIF5A, p97, and MYH2, showed that KIF5A had the highest sequence identity, at 24%. Notably, KIF5A contains histidines at positions equivalent to DFCP1 H195 and H323, whereas residues corresponding to R271 and T369 are absent (SI Appendix, Fig. S3). This suggests that KIF5A may also use a histidine-stack arrangement near the adenine base.

We next generated mutations in the histidine stack (H195 and H323), the potential gatekeeper residue T369 and the arginine finger R271 and analyzed their effect on nucleotide-dependent assembly. Interestingly, mutation of H195 and T369 had a strong detrimental effect on the expression level and protein purity of DFCP1, suggesting a key structural role of these residues which did not allow testing any mutants of these residues. H323 could be mutated and purified as an alanine, glutamate and arginine residue. H323A mutation completely abrogated nucleotide dependent dimerization of DFCP1 (Fig.2D), whereas H323E mutation did not affect the dimerization (Fig. 2E). Surprisingly, H323R mutation also did not show any shift of 0.3 ml between the ADP and AppNHp bound states, but the elution of both states was shifted towards the dimeric population, suggesting that this mutation can lock DFCP1 in a dimeric state independent of ADP/ AppNHp (Fig. 2F).

Furthermore, we tested the role of the R271 arginine finger with two mutations, alanine and glutamate. R271A mutation prohibited the AppNHp dependent shift of the elution peak, indicating that it blocks the dimerization (Fig. 2G), whereas R271E could still dimerize in the presence of AppNHp (Fig. 2H). This was surprising, as we expected R271E to be a dimerization deficient mutant. This suggests that a glutamate in this position can still form favorable interactions with the neighboring monomer in the presence of ATP.

We next quantified the ATPase activity of these mutants with help of ADP-Glo Kinase assay (Fig. 2I). All histidine-stack and arginine-finger mutants showed reduced ATPase activity compared to wild-type DFCP1. H323A and H323E were strongly impaired, whereas H323R retained partial activity. R271A showed reduced activity, while R271E was catalytically dead. Surprisingly, although H323E and R271E were found to be catalytically compromised, these mutations retain dimerization capability. This means that glutamate in these positions may be able to induce dimerization in the presence of AppNHp, but the interactions are catalytically irrelevant. These data show that both the histidine stack and the trans-acting arginine finger are required for efficient ATP hydrolysis.

We also searched for dimer interfaces outside the ATP binding pocket. We identified Q214 at the dimer interface, where the glutamine side chain reaches out in a surface pocket of the opposite monomer created by residues D270, Q325 and L272 (Fig. 2J and K). The Q214 side chain makes several H-bond interactions with the main chain carbonyl and amino groups of these residues. We therefore mutated Q214 to either arginine or glutamate to block these interactions. Both Q214R and Q214E mutants failed to show the clear AppNHp-dependent shift in size exclusion chromatography observed for wild-type DFCP1, indicating defective dimerization (Fig. 2L and M). In addition, these mutants abolished ATPase activity almost completely (Fig. 2N).

Together, these data show that DFCP1 ATPase activity depends on a composite active site formed by both monomers. The trans-acting arginine finger, the histidine stack, and Q214 at the dimer interface all contribute to nucleotide-dependent assembly and ATP hydrolysis. These results provide a structural explanation for how DFCP1 dimerization is a requirement for ATPase activity. At the same time, not all dimerizing mutants show ATPase activity as the dimerization needs to be catalytically relevant.

### DFCP1 dimer-interface mutants impair nucleotide-dependent clustering on PI3P-containing GUVs

To test how full-length DFCP1 organizes on PI3P-positive membranes, we used giant unilamellar vesicles (GUVs) prepared by PVA-gel hydration. GUV membranes were visualized with Atto 647N–DOPE, and recruitment of MBP-EGFP-DFCP1 was monitored by fluorescence microscopy. Wild-type DFCP1 was recruited to GUVs containing 0.1% PI3P, but not to control GUVs containing 0.3% PS, confirming that membrane binding depends on PI3P (Fig. 3A).

**Figure 3.**
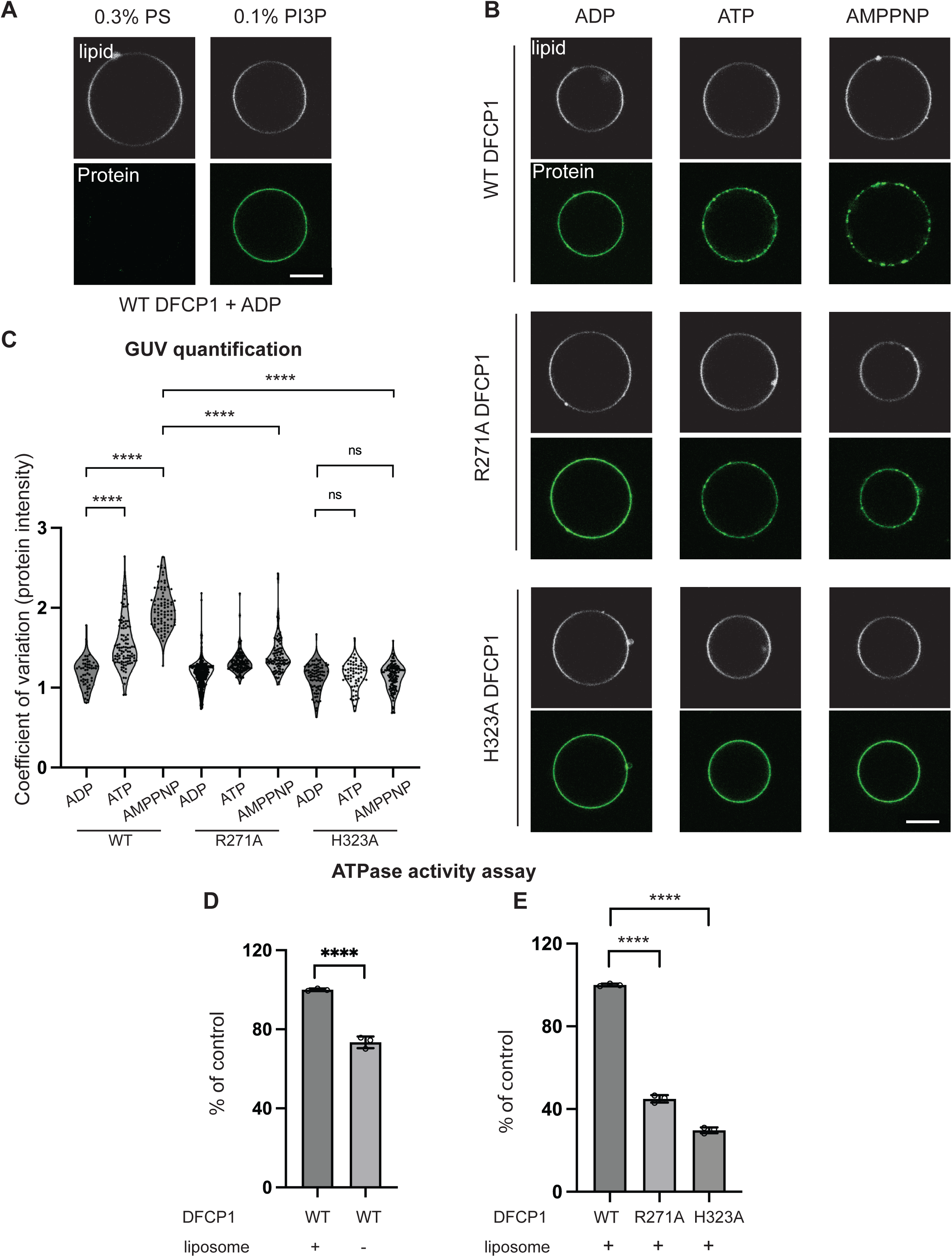
GUV localisation assay with full-length DFCP1 show clustering defect in DFCP1 ATPase mutants. (A) GUVs were prepared using the PVA-gel hydration method as described in *Materials and Methods*. Membrane recruitment of DFCP1 is dependent on PI3P. MBP-EGFP-DFCP1 is recruited to periphery of GUVs containing 0.1% PI3P. The membrane boundary was identified using 0.2% of the lipid dye Atto 647N–DOPE included in the lipid mixture. In contrast, no DFCP1 was observed on GUVs containing 0.3% POPS. (B) Effects of mutants on the dimer interface of DFCP1 on GUV membranes. Wild type DFCP1 shows puncta around the GUV membrane when incubated with ATP and AMPPNP. R271A mutant shows reduced puncta formation, while H323A mutant shows no puncta formation in the ATP and AMPPNP conditions. The coefficient of variation (CV = standard deviation / mean) of the protein intensity around each GUV perimeter is calculated as a measure of puncta formation. Quantification of GUVs showed significant puncta formation for wild-type DFCP1 at ATP (n=103 GUVs) and AMPPNP (n=87 GUVs) conditions, comparing to the ADP condition (n=55 GUVs). Both R271A (n=102 GUVs) and H323A (n=124 GUVs) mutants showed significant less puncta formation compared to the wild type in the AMPPNP condition. Circles in the violin plot represent independent data points. (C) ATPase activity of MBP-EGFP-DFCP1 is measured using Promega’s ADP-Glo TM kinase assay. Membrane increased the ATPase activity of full-length DFCP1. Both R271A and H323A mutants have reduced ATPase activity compared to the wild type. All results are from at least three independent experiments. P ≤ 0.0001 (****), ns = not significant. (Scale bar, 10 μm.)

We next asked whether nucleotide state affects the organization of membrane-bound DFCP1. In the presence of ADP, wild-type DFCP1 showed a relatively continuous distribution along the GUV membrane. In contrast, ATP and AppNHp, a non-hydrolysable ATP analogue, induced a more heterogeneous pattern, with DFCP1 accumulating in distinct puncta along the membrane (Fig. 3B). Wild-type DFCP1 showed significantly increased clustering under the ATP and AppNHp conditions compared with ADP (Fig. 3C). Thus, ATP-bound states promote clustering of DFCP1 on PI3P-containing membranes, presumably reflecting higher-order multimerization in the plane of the membrane.

We then tested whether residues at the DFCP1 dimer interface are required for this membrane organization. The R271A mutant, targeting the trans-acting arginine finger, was recruited to PI3P-containing GUVs but showed reduced clustering compared with wild-type DFCP1, particularly in the AppNHp condition. The H323A mutant, targeting the histidine stack, was also recruited to GUV membranes but failed to form pronounced puncta in either ATP or AppNHp (Fig. 3B). Quantification confirmed that both R271A and H323A showed significantly reduced clustering compared with wild-type DFCP1 in the AMPPNP condition, while H323A showed no significant nucleotide-dependent increase in clustering whatsoever (Fig. 3C).

We also measured DFCP1 ATPase activity in the presence and absence of liposomes. Liposomes stimulated ATPase activity of wild-type DFCP1, confirming our previous observations that membrane binding promotes DFCP1 catalytic activity (Fig. 3D, Nähse et al 2023). In the presence of liposomes, both R271A and H323A showed strongly reduced ATPase activity compared with wild-type DFCP1 (Fig. 3E). The reduction was more pronounced for H323A than for R271A, consistent with the stronger clustering defect observed on GUV membranes.

Together, these data show that PI3P recruits DFCP1 to membranes, while ATP-bound states promote DFCP1 clustering on the membrane surface. Mutations in the arginine finger or histidine stack impair both nucleotide-dependent clustering and membrane-stimulated ATPase activity. Thus, the DFCP1 dimer interface controls not only ATP hydrolysis, but also the higher-order organization of DFCP1 on PI3P-positive membranes.

### Loss of the DFCP1 ATPase domain alters omegasome-phagophore dynamics

To further investigate the role of the DFCP1 ATPase domain, we first asked whether DFCP1 lacking this domain can still localize to omegasomes, and whether it phenocopies ATPase-deficient DFCP1 mutants. For this, we expressed mNeonGreen-tagged DFCP1 lacking the N-terminal 417 amino acids, hereafter referred to as DFCP1 ΔATPase, in DFCP1 knockout U2OS cells (Fig. 4a). We then compared omegasome formation in DFCP1 knockout cells rescued with either wild-type DFCP1 or DFCP1 ΔATPase.

**Figure 4.**
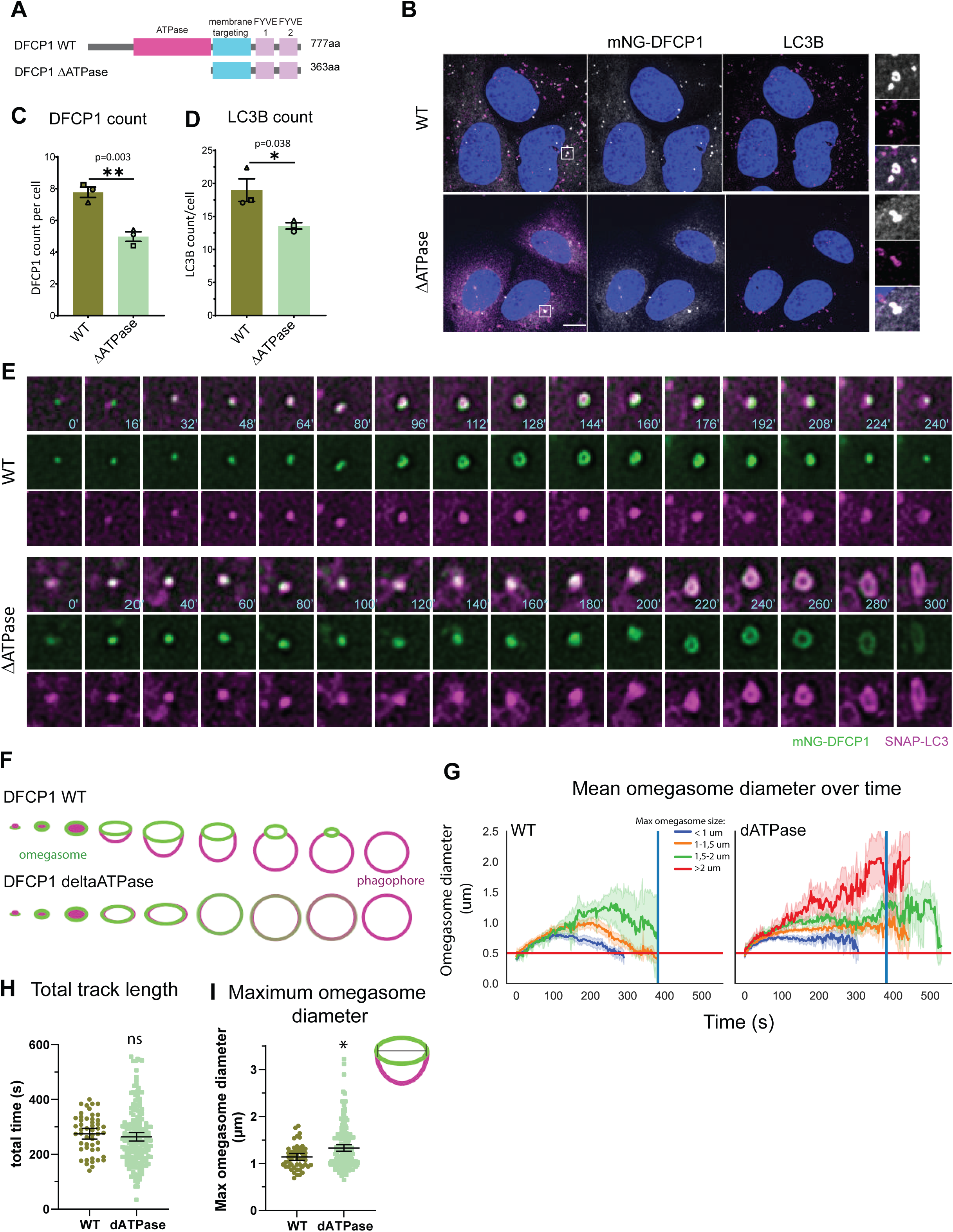
Loss of the DFCP1 ATPase domain alters omegasome-phagophore dynamics. (A) Schematic representation of wild-type DFCP1 and the DFCP1 ΔATPase construct. In DFCP1 ΔATPase, the N-terminal 417 amino acids containing the ATPase domain are deleted. (B) Representative confocal images of DFCP1 knockout U2OS cells rescued with mNG-tagged DFCP1 WT or DFCP1 ΔATPase. Cells were starved in EBSS for 2h, fixed, and stained for endogenous LC3B. Insets show magnified examples of DFCP1-positive omegasomes and associated LC3B-positive structures. Scale bars, 10 µm. Quantification of (C) DFCP1-positive puncta or (D) LC3B positive puncta per cell in DFCP1 knockout U2OS cells expressing mNG-DFCP1 WT or mNG-DFCP1 ΔATPase after 2h EBSS starvation. Data are shown as mean ± SEM. (E) Time-lapse microscopy of DFCP1 knockout U2OS cells co-expressing mNG-DFCP1 WT or mNG-DFCP1 ΔATPase together with SNAP-LC3. Time points are indicated in seconds. (F) Schematic model summarizing the altered omegasome–phagophore sequence in cells expressing DFCP1 ΔATPase. WT DFCP1-positive omegasomes expand, separate from the growing LC3-positive phagophore, and constrict. DFCP1 ΔATPase-positive omegasomes fail to constrict and instead display a fading phenotype, in which DFCP1 remains associated with the LC3-positive structure before disappearing. (G) Mean omegasome diameter over time in DFCP1 knockout U2OS cells expressing mNG-DFCP1 WT or mNG-DFCP1 ΔATPase. Tracks were grouped according to maximum omegasome diameter. (H) Quantification of total DFCP1 track length in cells expressing mNG-DFCP1 WT or mNG-DFCP1 ΔATPase. (I) Quantification of maximum omegasome diameter in cells expressing mNG-DFCP1 WT or mNG-DFCP1 ΔATPase.

Cells were starved in EBSS to induce both selective and non-selective autophagy, fixed, and the number of omegasomes per cell was quantified. Wild-type DFCP1-rescued cells formed on average approximately 8 omegasomes per cell. Surprisingly, DFCP1 ΔATPase was still able to localize to omegasomes. However, the number of omegasomes was significantly reduced compared to wild-type DFCP1 (Fig. 4B and C). Consistent with this, the number of LC3-positive structures was also reduced, indicating that phagophore production at omegasomes is impaired in cells expressing DFCP1 ΔATPase (Fig. 4D). This was further supported by a reduction in the number of WIPI2B-positive structures (SI Appendix, Fig. S4A), although the remaining WIPI2B structures were larger than in wild-type cells (SI Appendix, Fig. S4B).

We next performed live-cell imaging to analyse omegasome dynamics. Using DFCP1 knockout U2OS cells expressing mNG-tagged wild-type DFCP1 or DFCP1 ΔATPase together with SNAP-LC3, allowed us to follow both omegasome and phagophore formation over time. Both wild-type and ΔATPase cells form ring-like structures which expand. A striking difference was observed in the constriction behaviour of DFCP1 ΔATPase-positive omegasomes. In wild-type cells, DFCP1-positive omegasome rings initially expanded until the DFCP1 ring reached its maximum size, then the DFCP1 signal separated from the LC3-positive phagophore, which enlarged and extruded from the omegasome ring forming an autophagosome. Lastly, the omegasome ring constricted into a single DFCP1-positive dot. (Fig. 4E) (Nahse et al., 2023). Thus, at later stages, DFCP1 and LC3 occupied two distinct membrane structures.

DFCP1 ΔATPase-expressinig cells showed a different behaviour. While initial formation and expansion of the omegasome ring structure were highly similar to full-length DFCP1, DFCP1 ΔATPase-positive omegasomes failed to constrict upon reaching larger diameters (Fig. 4E and F). Moreover, in most DFCP1 ΔATPase-positive structures, the mNG-DFCP1 signal did not separate from the SNAP-LC3 signal. Although a phagophore still formed, it lacked the classical extrusion step observed in wild-type cells. Instead, the DFCP1 signal remained associated with the LC3-positive structure and gradually disappeared over time. We refer to this phenotype as “fading” (Fig. 4E and F).

Quantitative analysis over time confirmed these observations. Wild-type omegasomes initially expanded and subsequently constricted into a DFCP1-positive dot, with omegasome ring size correlating with constriction timing. In contrast, DFCP1 ΔATPase-positive rings failed to constrict back into a dot, irrespective of their maximum size (Fig. 4G). The overall duration of the DFCP1 localization was not significantly altered (Fig. 4H), but the maximum ring diameter was increased in DFCP1 ΔATPase-expressing cells, consistent with a complete loss of constriction (Fig. 4I). Moreover, approximately 75% of DFCP1 ΔATPase-positive omegasomes displayed the “fading” phenotype (Fig. 6A).

Together, these data show that the DFCP1 ATPase domain is dispensable for omegasome recruitment and ring formation, but is crucial for productive omegasome constriction. In its absence, omegasomes can still support phagophore formation, but maturation follows an altered sequence in which LC3-positive structures fail to extrude from the DFCP1 ring and the DFCP1 signal instead fades over time.

### Mutations in the DFCP1 dimerization interface cause abnormal omegasome phenotypes

To assess the importance of ATPase-domain dimerization for omegasome dynamics, we selected the R271A and H323A mutants for further analysis. These mutants were expressed in DFCP1 knockout U2OS cells, followed by EBSS starvation to induce autophagy and analyse omegasome formation.

Under fixed-cell conditions, we observed an increased number of DFCP1-positive puncta in cells expressing the arginine-finger mutant R271A, whereas cells expressing the H323A mutant showed a non-significant tendency towards reduced DFCP1 puncta formation (Fig. 5A and B).

**Figure 5.**
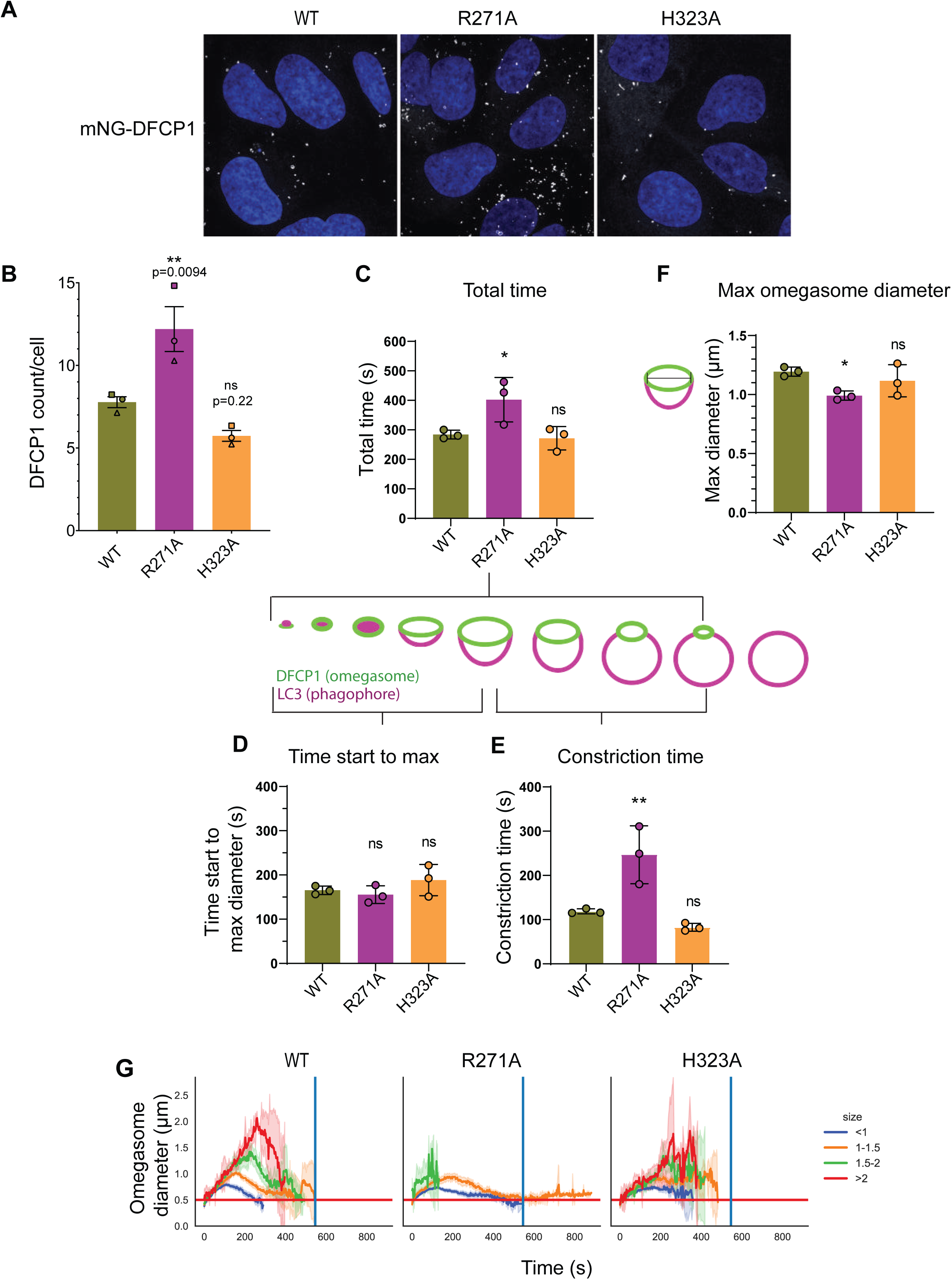
Omegasome dynamics are altered in arginine-finger and histidine-stack mutants. (A) Representative spinning-disk microscopy images of DFCP1-knockout U2OS cells expressing mNG-tagged DFCP1 WT, R271A, or H323A. Cells were starved in EBSS for 2 h, fixed, and imaged to assess DFCP1-positive omegasome formation. (B) Quantification of DFCP1-positive puncta per cell in cells treated as in A. (C–F) Quantification of DFCP1-positive omegasome tracks from live-cell imaging experiments. DFCP1-knockout U2OS cells expressing mNG-tagged DFCP1 WT, R271A, or H323A were starved in EBSS and subjected to time-lapse imaging. (C) Total track duration, measured from the first appearance of a DFCP1-positive structure to disappearance of the DFCP1 signal. (D) Time from first DFCP1 appearance to maximum omegasome diameter. (E) Constriction time, defined as the time from maximum omegasome diameter to disappearance of the DFCP1-positive structure. (F) Maximum omegasome diameter measured for individual tracks. (G) Omegasome diameter over time for DFCP1 WT and the indicated mutants, grouped according to maximum omegasome diameter.

We next performed live-cell imaging to determine how these mutations affect omegasome dynamics (SI Appendix, Fig. S5A). The R271A mutant required more time to complete the omegasome sequence, whereas the overall timing was not significantly altered for the other mutants (Fig. 5C). More detailed analysis showed that the time from DFCP1 appearance to formation of the omegasome ring was not significantly affected in any of the three mutants. However, the constriction phase, defined as the time from maximum ring diameter to disappearance of the DFCP1-positive dot, was strongly delayed in R271A-expressing cells, taking approximately twice as long as in wild-type cells. This phenocopies our previously identified T189A and K193A mutants, defective in ATP hydrolysis and binding, respectively, which similarly displayed delayed constriction (Nahse et al., 2023). Like these mutants, R271A did not alter the overall omegasome morphology: omegasomes still progressed through initiation, expansion, and eventual constriction into a single DFCP1-positive dot, albeit with delayed constriction.

While analysing individual live-cell tracks, we also noticed clear morphological differences between the mutants (SI Appendix, Fig. S5A). The maximum omegasome diameter was reduced in R271A-expressing cells (Fig. 5F and G), but R271A-positive omegasomes progressed through all canonical phases (SI Appendix, Fig. S5A; Fig. 6B). In contrast, analysis of H323A-positive omegasomes revealed a phenotype resembling that of DFCP1 ΔATPase. In approximately 60% of H323A-positive omegasomes, the DFCP1 signal failed to separate from the LC3 signal (Fig. 6A). Instead, these omegasomes displayed the previously described “fading” phenotype (Fig. 3F), in which the DFCP1 signal remained associated with the LC3-positive structure and gradually disappeared over time (Fig. 6A).

**Figure 6.**
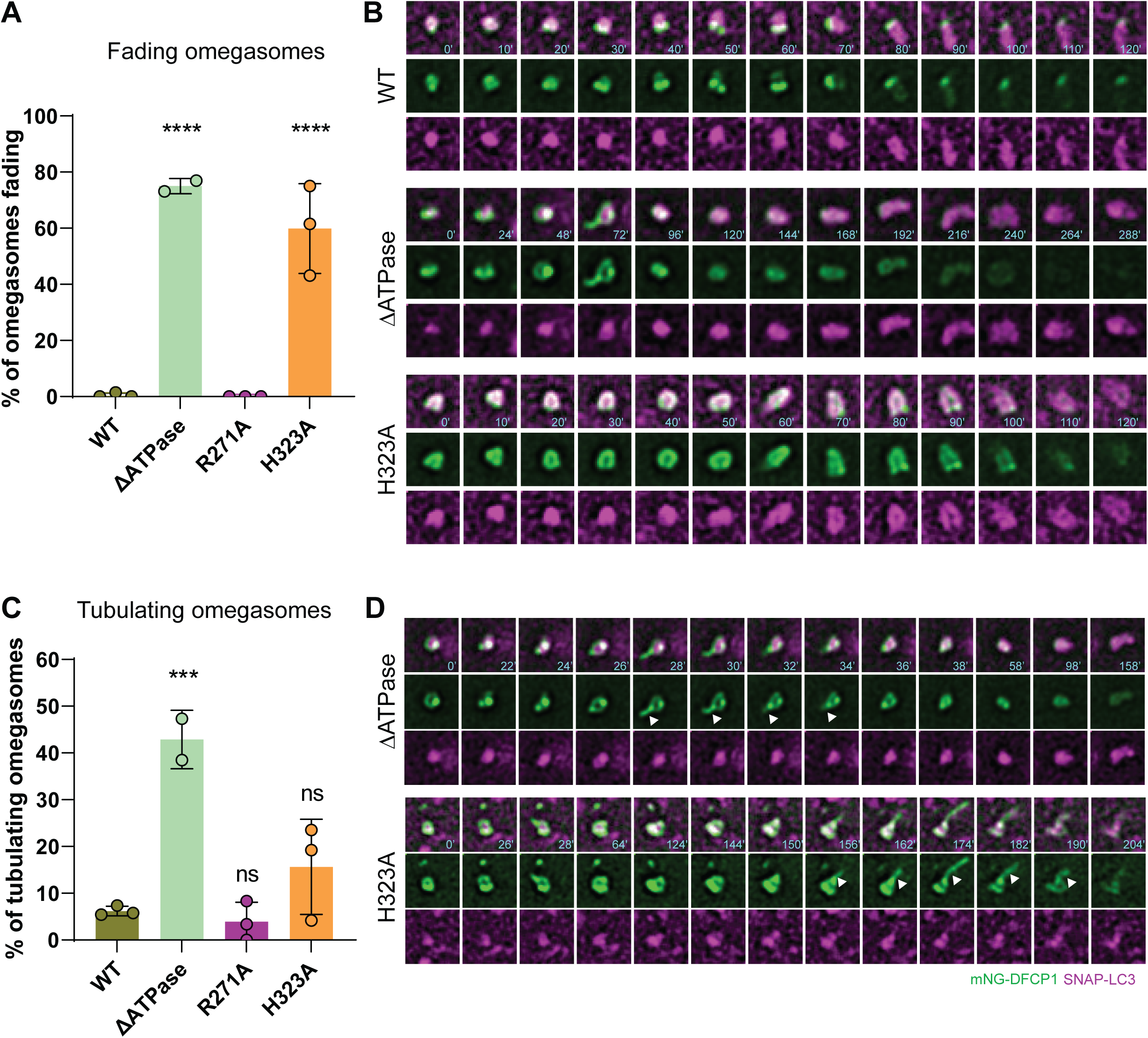
Morphological changes of omegasomes in DFCP1 arginine-finger and histidine-stack mutants. DFCP1-knockout U2OS cells expressing mNG-tagged DFCP1 WT, DFCP1 ΔATPase, or the indicated point mutants together with SNAP-LC3 were starved in EBSS and analysed by live-cell spinning-disk microscopy. Time-lapse sequences show mNG-DFCP1 and SNAP-LC3 signals, with time indicated in seconds. Quantifications show the percentage of omegasomes displaying the indicated phenotype. Data are shown as mean ± SEM from 2–3 independent experiments. (A) Quantification of fading omegasomes. Fading was defined as an omegasome event in which the mNG-DFCP1 signal remained associated with the SNAP-LC3-positive structure and gradually disappeared over time, without clear separation of the DFCP1-positive omegasome from the LC3-positive phagophore or constriction into a single DFCP1-positive dot. (B) Representative time-lapse sequences of fading omegasomes in cells expressing DFCP1 WT, ΔATPase, or H323A. (C) Quantification of tubulating omegasomes. Tubulating omegasomes were defined as DFCP1-positive omegasomes that formed transient DFCP1-positive tubular extensions during the imaging sequence. These tubules extended from the omegasome and retracted or disappeared within seconds. (D) Representative time-lapse sequences of tubulating omegasomes in cells expressing DFCP1 ΔATPase or H323A.

H323A and DFCP1 ΔATPase-positive omegasomes also shared an additional feature. During ring formation and expansion, DFCP1-positive tubules frequently emerged and retracted within seconds, indicating highly dynamic membrane remodeling. This phenotype was particularly prominent in the ΔATPase mutant, where more than 40% of omegasomes formed DFCP1-positive tubules (Fig. 6C). This suggests that DFCP1 can promote membrane deformation independently of ATP hydrolysis, possibly through clustering on PI3P-positive membranes.

Together, these data show that mutations in the DFCP1 dimerization interface impair omegasome dynamics but do so in distinct ways. R271A primarily delays constriction without disrupting the canonical omegasome sequence, whereas H323A phenocopies DFCP1 ΔATPase by causing a fading phenotype. These findings support a model in which ATPase-dependent dimerization is required not only for omegasome constriction, but also for coordinating the morphology and sequence of omegasome–phagophore dynamics.

## Discussion

Our crystal structures of the DFCP1 ATPase domain bound to ADP or the non-hydrolysable ATP analogue AppNHp reveal a composite active site formed at the dimer interface. This active site contains a Walker-A-like phosphate-binding loop, a trans-acting arginine finger, and a histidine sandwich that positions and orients the nucleotide between opposing protomers. Structure-guided mutagenesis, biochemical assays, GUV-based membrane reconstitution, and live-cell imaging together show that these elements are required not only for ATP hydrolysis and multimerization, but also for the correct timing and morphology of omegasome progression in cells.

To assess evolutionary conservation within DFCP1, we aligned the top 150 homologous sequences across organisms using the ConSurf server (Ashkenazy et al., 2016) (SI appendix, Fig. S6A-B). Interestingly, the N-terminal domain of DFCP1 appears variable except a few residues, however, ATPase domain followed by the ER-targeting domains are the most conserved elements in DFCP1. The Zn^2+^ coordinating and the PI(3)P binding residues also show high conservation in the two FYVE domains. The strong conservation of ATPase, ER-targeting domain as well as the linker connecting the two modules, indicates that DFCP1 might follow an ATP-hydrolysis driven conformational switch mechanism as seen in several Dynamin family membrane remodelling proteins. In the ATPase domain, H195 and H323 are highly conserved, although in some organisms they are replaced by tyrosine, phenylalanine, or aspartate. These aromatic substitutions could still allow π–π stacking with the adenine base. H195 is more strongly conserved than H323, which may explain why substitutions at position 195 were difficult to generate, as mutations at this site may compromise structural integrity. R271 and T369 are highly conserved residues underlining their critical role in ATP binding and hydrolysis. Mapping conservation scores onto the AlphaFold3 prediction (Abramson et al., 2024) of the DFCP1 ATPase domain dimer in complex with ATP and Mg^2+^ shows that the ATP binding site and the dimeric interface is highly conserved (SI appendix, Fig. S6C-E). We propose that H195, H323, and T369 contribute to DFCP1 selectivity for ATP whereas R271 is crucial for ATP hydrolysis. In addition, there might be several secondary interfaces as shown for Q214 residue that stabilize the dimeric interface outside the ATP-induced primary interface.

R266 in the DFCP1 ATPase domain has been reported to be mutated to glutamine in patients with endometrial and colorectal cancer (Ismail et al., 2023). This residue is also highly conserved across organisms. In our crystal structures, R266 forms a hydrogen bond with E320 and lies in close proximity to the adenine base of ATP. The R266Q mutation has been reported to reduce DFCP1 ATPase activity, which our structures suggest may result from destabilization of the adenine-base binding pocket.

Among the point mutants, DFCP1 H323A most closely resembled the phenotype observed upon complete deletion of the ATPase domain. H323A-positive omegasomes frequently failed to separate from LC3-positive structures and instead displayed the “fading” phenotype, defined by persistent colocalization of DFCP1 with LC3 followed by gradual loss of the DFCP1 signal, without the normal separation and constriction steps. This was unexpected because both H323A and R271A disrupt nucleotide-dependent dimerization and reduce ATPase activity, yet they produced different cellular phenotypes. One explanation could be that even though R271, together with T189 and K193 are catalytically important residues, they could still be compensated by another residue if enough time is provided, whereas H323 being only one of the three residues (H195, T369 and H323) that coordinate with adenine base is more difficult to be compensated and results into defective ATP recruitment and dimerization. Where H323 is crucial for binding both ATP and ADP, R271 is more critical for γ-phosphate binding and thus regulates conformational switch followed by hydrolysis. Therefore, presence of H323 is a prerequisite for recruitment of R271.

We set out to address a central question in DFCP1 biology: how ATP binding and hydrolysis contributes to omegasome formation. We show that ATP hydrolysis does not control membrane recruitment, which instead is determined by the C-terminal half of DFCP1 together with the ER-targeting and FYVE domains. Instead, our crystal structures, biochemical assays, membrane-reconstitution experiments, and cell-imaging analyses identify ATP-dependent dimerization as a key mechanism that promotes local DFCP1 clustering and omegasome structure. These findings provide a framework for future studies of full-length DFCP1 at membranes and for understanding how ATP hydrolysis is coupled to membrane remodelling.

## Materials and Methods

### Cloning and site-directed mutagenesis

DFCP1 construct corresponding to residues 146-416 was cloned into pET22b vector using NdeI and XhoI restriction sites. For crystallization, we cloned DFCP1 146-409 construct into pET28a(+)-TEV vector to enable tag removal. For expression in *Sf*9 insect cells (*Spodoptera frugiperda,* TriEX^TM^ *Sf*9 cells, Novagen) DFCP1_146-409_ construct was cloned into baculovirus expression/transfer vector pFB-6HZB vector that comprises a N-terminal His_6_-and Z-basic tag followed by a TEV protease cleavage site. Site-directed mutagenesis was performed by PCR using primers designed to introduce mutation. Following amplification, DpnI was added to each reaction mixture to digest the methylated parental plasmid template. The reactions were incubated overnight at 37 °C to ensure complete digestion of the template DNA. After reaction clean up, the plasmid was transformed and the mutation was confirmed by sanger sequencing of single colonies.

### Bacterial and insect cell culture

For recombinant protein expression, plasmids were transformed into chemically competent *E. coli* BL21 cells. Single colonies were picked from selective LB agar plates and used to inoculate LB medium for secondary culture, supplemented with the appropriate antibiotic. Cultures were grown overnight at 37 °C with shaking at 180 rpm. Secondary bacterial cultures for recombinant protein expression were inoculated from primary overnight cultures. Secondary cultures were inoculated to 0.05 OD_600_ units from primary cultures. Cultures were incubated at 37 °C with shaking until reaching mid-logarithmic growth phase (OD600 ≈ 0.6–1.0). Protein expression was induced by the addition of 0.4 mM isopropyl β-D-1-thiogalactopyranoside (IPTG). Following induction, cultures were incubated in shakers at 18 °C overnight. Cells were harvested by centrifugation at 4500 rpm, 4 °C, for 30 min, and the resulting pellets were resuspended in lysis buffer (50 mM Tris-HCl pH-7.5, 500 mM NaCl, 25 mM imidazole, 0.5 mM TCEP, 0.1 % lysozyme, 2.5 mU/ml DNAse) containing 25% glycerol and flash frozen and stored at −80 °C for further purification.

For insect cell expression, exponentially growing *Sf*9 cells (2e^6^ cells/mL in Insect-XPRESS medium, Lonza) were infected with high-titer baculovirus suspension. After 66 h of incubation (27 °C and 90 rpm), cells were harvested by centrifugation.

### Protein Purification

The bacterial pellets were thawed and subsequently lysed by probe sonication using a pulsed program of 1 s ON and 1 s OFF at 40% amplitude for a total sonication time of 15 min. Samples were kept cold during sonication to prevent overheating. Following cell disruption, insoluble fraction was removed by centrifugation at 18,000 rpm for 45 min at 4 °C. The clarified supernatant containing soluble proteins was collected and used for subsequent affinity purification. With the help of His-tag, proteins were purified using Ni²⁺ affinity chromatography. Ni-NTA agarose beads were first added to a gravity-flow bench column and washed with Milli-Q water to remove the storage solution (20% Ethanol). The resin was subsequently equilibrated with lysis buffer. The lysate was then added to the equilibrated resin and gently resuspended. The lysate–resin mixture was transferred into a bottle and incubated on a bench roller at 18 rpm for 1 h at room temperature to allow binding of His-tagged proteins to the Ni-NTA resin. After incubation, the lysate–resin mixture was returned to the gravity-flow column, allowing the unbound fraction to pass through. The resin was subsequently washed twice with lysis buffer to remove non-specifically bound proteins. Bound proteins were eluted using elution buffer (50 mM Tris-HCl pH-7.5, 500 mM NaCl, 250 mM imidazole, protease inhibitor cocktail and 25% glycerol)), and the eluate was collected and analyzed by SDS-PAGE to assess protein purity. Fractions containing the protein of interest were pooled and subsequently concentrated using Amicon centrifugal filters for size exclusion chromatography.

All pET22b-derived DFCP1 constructs were loaded onto a HiLoad 16/600 Superdex 75 pg column. Samples were injected via a sample loop and separated under isocratic elution conditions in the gel filtration buffer (50 mM Tris-HCl pH-7.5, 200 mM NaCl and 1 mM TCEP). Protein elution was followed by UV absorbance at 280 nm and 254 nm, and fractions from the monodisperse peak were pooled for downstream analysis. Protein aliquots were therefore stored at −80 °C and thawed only when required for experiments.

The full-length, codon-optimized DFCP1 gene was synthesized (Twist Bioscience) and cloned into a pCAG vector. The DFCP1 construct includes sequence for a TEV-cleavable MBP tag at the N terminus, followed by sequence for a EGFP tag, and a 3C protease cleavage site. A total of 1 mg plasmid DNA and 4 mg PEI (Sigma-Aldrich) were used to transfect HEK293F GnTI-cells at a density of 1.8 × 106 cells per ml per l. The cells were pelleted after 72 h of transfection and lysed in lysis buffer (50 mM HEPES, pH 7.4, 300 mM NaCl, 1mM MgCl_2_, 1mM TCEP, 10% glycerol, 1% Triton X-100 and protease inhibitor). The cleared supernatant after centrifugation was applied to Amylose Resin (NEB) and incubated for 2 h. The resin was washed with wash buffer (50 mM HEPES, pH 7.4, 300 mM NaCl, 1mM MgCl_2_, 1mM TCEP) first with 1% Triton X-100 and then without Triton X-100. The complex was eluted by elution buffer (50 mM HEPES, pH 7.4, 150 mM NaCl, 1mM MgCl_2_, 1mM TCEP, 100mM maltose) and incubated with 100 uM ATP overnight at 4 °C. Further purification was performed by SEC using a Superdex 200 10/300 GL column equilibrated with the elution buffer but without maltose. The fractions corresponding to monomeric DFCP1 were pooled and concentrated.

### DFCP1-146-409 crystallization construct purification

DFCP1_146-409_ that was expressed in *Sf*9 insect cells contained an N-terminal His_6_-Z-tag and a TEV protease cleavage site. *Sf*9 cells were pelleted and washed with PBS, following resuspension in *Sf*9 lysis buffer (50 mM HEPES pH 7.4, 500 mM NaCl, 20 mM imidazole, 0.5 mM TCEP, 5% glycerol) and lysis by sonication. The lysate was cleared by centrifugation and loaded onto a NiNTA column. After vigorous rinsing with *Sf*9 lysis buffer, the His_6_-Z-tagged protein was eluted in *Sf*9 lysis buffer containing 300 mM imidazole. Immediately thereafter, the eluate was diluted to 250 mM NaCl concentration with buffer containing no NaCl and loaded on an HiTrap^TM^ SP FF (Cytiva). His_6_-Z-TEV-DFCP1 was eluted with a gradient ranging from 250 mM to 2.5 M NaCl and treated with TEV protease overnight to cleave the His_6_-Z-tag. The cleaved tag, uncleaved protein and TEV protease were removed by another combined SP sepharose NiNTA step. Finally, the protein was concentrated and subjected to GF in storage buffer (20 mM HEPES pH 7.4, 150 mM NaCl, 0.5 mM TCEP, 5% glycerol) using an Äkta FPLC system combined with a HiLoad^TM^ 16/600 Superdex^TM^ 75 pg (Cytiva). Determination of protein concentration was carried out by absorption measurements at λ = 280 nm (Nanodrop) using molecular weights and theoretical extinction coefficients. Protein samples were aliquoted, flash-frozen in liquid nitrogen and stored at −80°C until use.

*E. coli* BL21(DE3) lysate for the 146-409 pET28a(+)-TEV construct was first subjected to Ni²⁺ affinity chromatography using a gravity-flow column. The elution fractions containing the target protein were pooled and incubated with TEV-His_6_ protease to remove the His_6_ tag and dialyzed overnight at 4 °C using dialysis buffer (50 mM Tris-HCl pH-7.5, 150 mM NaCl, 0.5 mM EDTA, 1 mM TCEP), followed by an additional 1h incubation at room temperature. After dialysis, the samples were centrifuged at 14,000 rpm at 4°C to remove aggregates. To separate cleaved from uncleaved DFCP1-146–409-TEV-His_6_ and TEV-His_6_ protease, the digested sample was loaded onto a HiLoad 16/600 Superdex 75 pg column connected in series to a His-Trap FF 5 mL column in the presence of an additional 25 mM imidazole in the gel filtration buffer. Finally, a third SEC run (Superdex 10/300 GL increase 75pg) was performed as a polishing step to further improve sample homogeneity for downstream crystallization attempt.

### Protein crystallography and Structure refinement

Tag-free DFCP1-146-409 was crystallized at the concentration of 12 mg/ml in the presence of either 5 mM AppNHp or 5mM ADP.Mg and 10 mM MgCl_2_. Protein crystallization was performed with the help of Mosquito liquid handling robot at RT. Crystallization screening was performed at 18°C with the help of commercial coarse screens with 300 nL of protein and 300 nL of mother liquor solution and sitting drop vapor diffusion method. DFCP1-ADP.Mg crystals were observed in the mother liquor containing 0.2 M ammonium sulfate, 25% w/v PEG 3350 and 0.1 M HEPES pH-7.5. Another crystal of DFCP1-ADP.Mg was obtained in the mother liquor containing 0.2 M ammonium sulfate, 0.1 M Bis-Tris pH-6.5, 25% w/v PEG 3350. DFCP1-AppNHp complex crystals were obtained in the mother liquor containing 0.2 M magnesium chloride hexahydrate, 0.1 M Tris pH 8.5, 25 % w/v PEG 3350. Crystals were flash-frozen in liquid nitrogen using a mother liquor supplemented with 25% ethylene glycol as a cryoprotectant.

### Data acquisition, structure solution and refinement

The diffraction data was collected at synchrotron X-ray source and 80 K temperature. DFCP1-ADP.Mg crystals diffracted to 1.9 Å resolution whereas the DFCP1-AppNHp complex diffracted to 2.3 Å resolution. Xia2 and XDS was used to do the autoprocessing and data reduction. Data scaling, integration and merging was carried out with Aimless (Evans, 2011) in the CCP4 software suite 9.0.006 (Agirre et al., 2023) ADP bound state was crystallized in monoclinic (P 1 21 1) space group whereas AppNHp complex crystals were found in orthorhombic (P 21 2 21) space group. We got another diffraction data for ADP bound crystals which were packed in C2 2 21 space group at the resolution of 2.4 Å. Alphafold3 model of DFCP1 was generated using the sequence of the corresponding constructs. Phaser (CCP4 suite) was used to do molecular replacement. The modeling was carried out with the help of Coot 0.9.8.95 and phenix.refine refinement program (Phenix 1.21.2-5419) (Adams et al., 2010). The data collection and refinement statistics are summarized in **Table 1**. Validation of the final models was performed using MolProbity (Chen et al., 2010). The resulting model was visualized with Pymol 3.1.3.1 (Schrödinger).

### Nucleotide-dependent dimerization determination with gel filtration

Nucleotide-dependent dimerization of the DFCP1 ATPase domain (DFCP1-146–416-His_6_ pET22b) and selected mutants was assessed by analytical size exclusion chromatography with ADP or AppNHp incubations. Protein was thawed from the freezer in an ice box and then centrifuged to remove aggregates from the sample at 14,000 rpm and 4°C. The concentration was determined by UV/Vis spectroscopy and adjusted to 5 mg/mL. For each construct, two aliquots were prepared and incubated overnight with either 5 mM ADP or 5 mM AppNHp (adenosine 5′-(β,γ-imido)triphosphate) to allow equilibration of nucleotide-bound states prior to chromatography. Before injecting the samples into the FPLC, the samples were centrifuged again at 14,000 rpm and 4°C.

Gel filtration runs were performed with Superdex 75 Increase 10/300 GL column. The column was equilibrated, and samples were applied at a flow rate of 0.50 mL/min, followed by isocratic elution at 0.25 mL/min for 1.20 column volume. Gel filtration buffer (50 mM Tris-HCl pH-7.5, 1 mM TCEP) was used containing low salt (50 mM) and 10 mM MgCl_2_ as additive. Elution was monitored by UV absorbance at 280 nm (UV1), 254 nm (UV2) and 214 nm (UV3). Chromatograms were exported from the ÄKTA system and processed in OriginPro for plotting and determination of peak elution volumes.

### ADP-Glo™ Kinase Assay

ATPase activity of DFCP1 variants was quantified using the ADP-Glo™ assay (Promega), a coupled luminescence-based assay that measures ADP generated during ATP hydrolysis. All reactions were performed in a White Corning® 384-well microplate (CLS4512) with a final reaction volume of 5.0 µL per well using the assay buffer (50 mM Tris-HCl pH-7.5, 50 mM NaCl, 20 mM MgCl_2_,0.1 % glycerol and 0.01 % Triton X-100). Nanolitre volumes of protein and ATP were dispensed using an Echo acoustic liquid handler, and buffer volumes were adjusted accordingly to maintain a constant final volume.

For ATPase measurements, the protein concentration was varied while the ATP concentration was held constant. Final DFCP1 concentrations were 9.0 µM, 8.0 µM, 7.0 µM, 6.0 µM, 5.0 µM, 4.0 µM, 3.0 µM, 2.0 µM, 1.0 µM, 0.5 µM, and 0.25 µM, while ATP was kept constant at 0.5 mM. For this setup, a corresponding 0.5 mM ADP standard curve was used. Negative controls consisted of wells containing buffer and ATP but no protein. After plate setup, the plates were briefly centrifuged and incubated for 20 min at room temperature. Reactions were then processed according to the manufacturer’s instructions by addition of ADP-Glo™ reagent to terminate the reaction and deplete residual ATP, followed by addition of Kinase Detection Reagent to convert ADP to ATP and generate a luciferase/luciferin-based luminescence signal. After the final incubation step, luminescence was recorded on a PHERAstar FSX plate reader in luminescence mode using a gain setting of 1800.

For data analysis, luminescence values were first blank-corrected and then converted into ADP concentrations (µM) using the corresponding ADP standard curve equation.

Reaction rates were calculated by dividing the amount of ADP formed by the reaction time (30 min) and are reported as µM ADP/min. For graphical comparison of ATPase activity, linear regression was applied to the reaction rate versus ATP concentration data, and the resulting slopes were normalized to the WT reference for the generation of the bar plots.

### GUV Formation

GUVs containing either 0.1 mol% PI3P or 0.3 mol % POPS, and DOPC (either 99.7 mol % or 99.5 mol %), the lipid fluorophore Atto 647N DOPE (0.2 mol%) were prepared in 361 mOsm sucrose using the polyvinyl alcohol (PVA)-gel hydration-based method as in Weinberger et al. (ref). In Brief, lipids were mixed in chloroform at a total concentration of 1.8 mM. 15 μL solution of the lipid mixture was spread on a 5% wt/vol PVA film dried on a 22 mm circular coverslip (Hampton Research) and then put under vacuum for at least 2 h to form a dry lipid film. 150 μL of 361 mOsm sucrose solution was used to hydrate the dried lipid film for 0.5 h at room temperature to produce GUV dispersion and then was collected and stored at 4 °C and used within 48 hours.

### Protein-GUV interaction and Confocal Microscopy

The imaging chamber (Lab-Tek II chambered cover glass, Fisher Scientific) was incubated with a 5 mg/mL solution of BSA for 30 min and washed three times with the reaction buffer (50 mM HEPES at pH 7.4, 150 mM NaCl, and 1 mM TCEP, 1mM MgCl_2_, 361 mOsm).

DFCP1 with different nucleotides were first mixed at a volume of 195 μL in a microcentrifuge tube at room temperature and immediately transferred into the imaging chamber. A total of 5 μL of GUVs was then added to the chamber. After a 10-min incubation, images were acquired on a Nikon A1 confocal microscope with a 63× Plan Apochromat 1.4 numerical aperture objective. Identical laser power and gain settings were used for each set of conditions. The reconstitution experiments were performed at room temperature.

### Liposome formation

A lipid mixture was prepared in a glass vial using the following lipid composition: 84.8 mol %, 10 mol % cholesterol, 5 mol % PI3P, and 0.2 mol % DiR. To form a thin film on the glass wall, the glass vial was slowly shaken on a vortex while drying under nitrogen gas. The glass vial was then placed in a vacuum oven overnight at room temperature to evaporate any remaining solvent. Lipids were hydrated in a lipid buffer (50 mM HEPES, pH 7.4, 150 mM NaCl) to a final concentration of 1.8 mM for 1 h, with intermittent vortexing during hydration. The solution was transferred to a 15-ml Eppendorf tube and subjected to nine freeze–thaw cycles using liquid nitrogen and a 40 °C water bath. The lipid mixture was either stored at −80 °C or immediately extruded using an Avanti Polar Lipids Mini Extruder (610023) at least 41 times through a 200-nm filter (Whatman Nuclepore Track-Etched Membranes, diameter of 19 mm) for DFCP1 kinase activity assays.

### ATPase Assay with liposomes

All enzyme activity by measured by the generation of ADP using Promega’s ADP-Glo TM Kinase Assay. All enzymes were assayed at 2 μM (DFCP1 wide type and mutans) and 0.2 mM liposome in 20 mM HEPES, pH 8.0, 100 mM NaCl, 1 mM ATP, and 10 mM MgCl 2 base buffer. Reactions were allowed to proceed at 37 °C for 1 hour before being quenched by the addition of Promega ADP-Glo TM Reagent at a 1:1 ratio. Assay conditions follow the recommended timing and ratios found in the Promega ADP-Glo TM Kinase Assay manual. The reaction + ADP-Glo TM Reagent solution was incubated for 40 minutes before the addition of Promega Kinase Detection Reagent at a 1:1 ratio (reaction + ADP-Glo TM Reagent: Kinase Detection Reagent) and incubated for 30 minutes prior to measuring reaction luminescence.

### Image Analysis GUVs

Quantification of fluorescence signal distribution along GUV membranes was performed using a custom Fiji (ImageJ) macro script (included in the supporting information). Membrane channel images are processed to identify individual GUVs based on contrast enhancement and auto thresholding, filtering by area and circularity to exclude irregular or fragmented objects. For each GUV, a binary mask is created and optionally shrunk inward by a defined number of pixels to restrict analysis to the membrane region. Using this mask, a ring region of interest (ROI) is generated corresponding to the membrane, and the mean and standard deviation of protein intensity within the ring are measured after background subtraction. The coefficient of variation (CV = standard deviation / mean) of the protein signal around each vesicle perimeter is calculated as a measure of fluorescence heterogeneity. For each image, the macro outputs a summary table including GUV area, mean membrane intensity, mean signal intensity, and calculated CV values. Results are compiled into a single spreadsheet and overlay images showing detected vesicles and analysis rings are optionally saved for quality control and manual inspection.

### Statistical Analysis GUVs

Statistical analysis was performed with GraphPad Prism 9.0. The data of the GUV-binding and ATPase activity assays were analysed by Student’s t test and one-way analysis of variance. Significance between the two calculated areas was determined using a Student’s two-tailed unpaired t test.

#### Antibodies

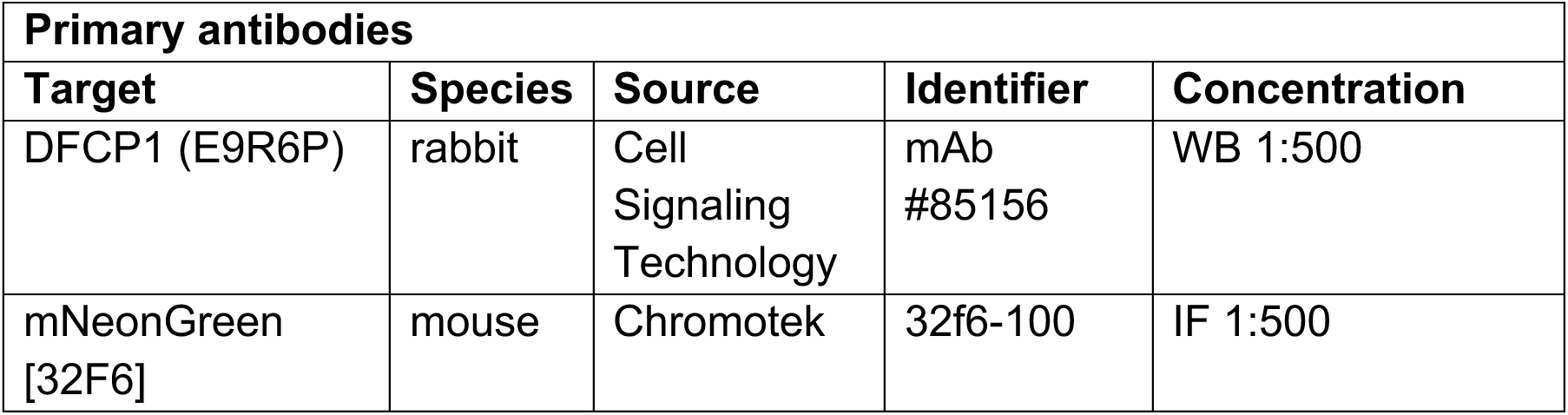
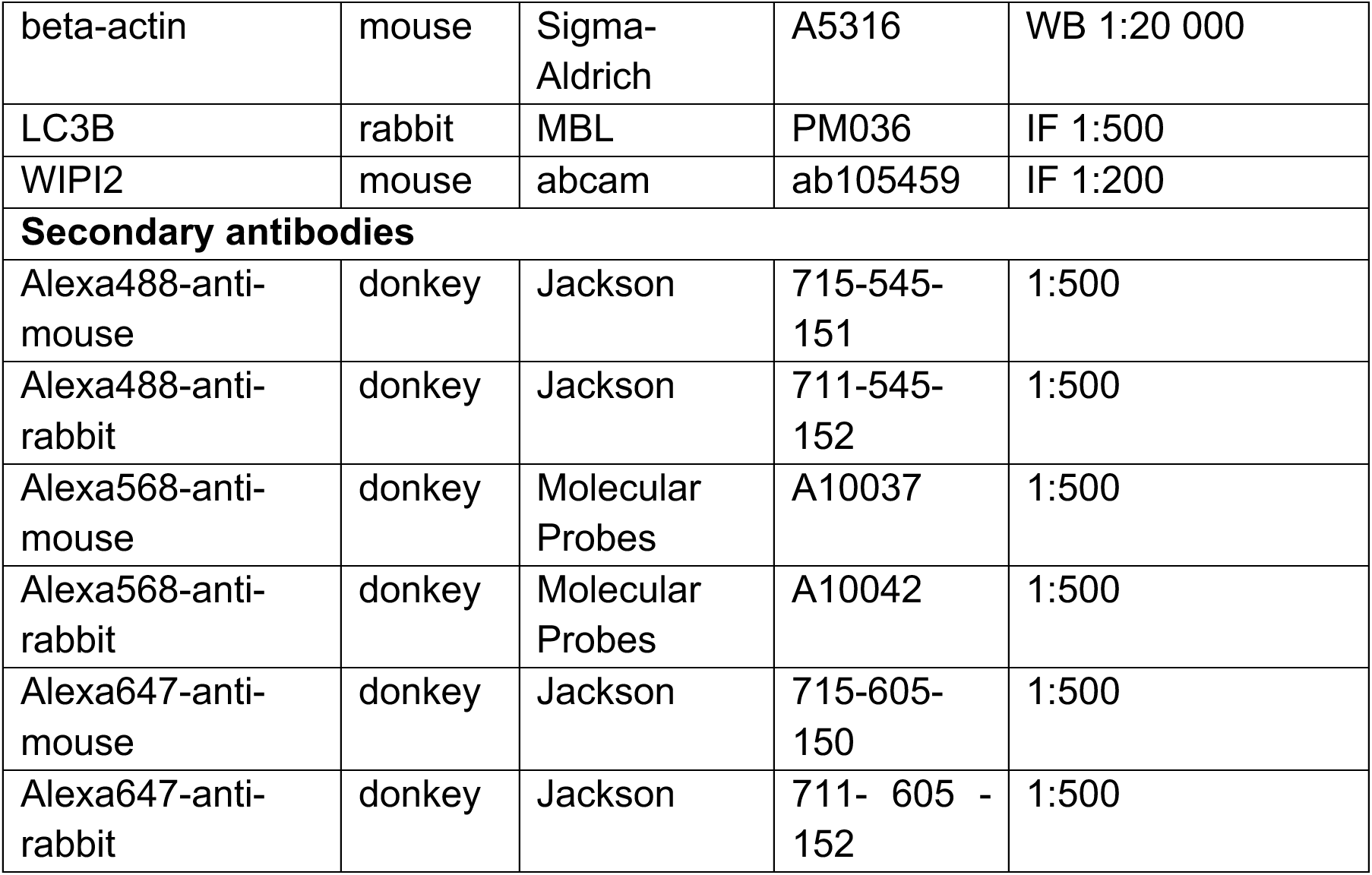

#### Constructs

|  |  |
| --- | --- |
| pCDH-PGK_mNeonGreen-DFCP1_IRES_Puro | (Nahse et al., 2023) |
| pCDH-PGK_mNeonGreen-DFCP1_R271A_IRES_Puro | this study |
| pCDH-PGK_mNeonGreen-DFCP1_H323A_IRES_Puro | this study |
| pCDH-PGK_mNeonGreen-DFCP1_H195A_IRES_Puro | this study |
| pCDH-PGK_SNAP_LC3B_IRES_Neo | (Nahse et al., 2023) |

#### Cell culture

U2OS cells (ATCC: HTB-96) were maintained in DMEM (D0819; Sigma-Aldrich) supplemented with 10% FCS (F7524; Sigma-Aldrich), 5 U/ml penicillin and 50 μg/ml streptomycin at 37°C with 5% CO_2_. hTERT-RPE-1 cells (ATCC: CRL-4000) were grown in DMEM/F12 medium (31331–028; Gibco-BRL) with 10% FCS, 5 U/ml penicillin and 50 µg/ml streptomycin at 37°C with 5% CO_2_. Parental cell lines and its derivatives were regularly tested for mycoplasma. For starvation experiments, cells were washed twice with EBSS (24010-043; GIBCO BRL) and incubated in EBSS for the indicated time.

### Generation of cell lines

DFCP1-knockout U2OS cells generated previously (Nahse et al., 2023) were used to generate rescue cell lines expressing DFCP1 WT or the indicated mutants. Stable rescue lines were generated by lentiviral transduction using a third-generation lentiviral vector system, as previously described (Campeau et al., 2009). All constructs were expressed from a phosphoglycerate kinase (PGK) promoter. Following transduction, cells were selected for stable integration of the expression cassette using puromycin (1 μg/mL) or geneticin (500 μg/mL).

### Immunoblotting

Cells were washed with ice-cold PBS and lysed in 2.8x Laemmli Sample buffer (BioRad) containing 200 mM DTT. Cell lysates were subjected to SDS-PAGE on a 4–20 % (567–1094; Bio-Rad) gradient gel and blotted onto PVDF membranes (BioRad). Membranes incubated with fluorescently labelled secondary antibodies (IRDye680 and IRDye800; LI-COR) were developed by Odyssey infrared scanner (LI-COR). Membranes detected with HRP labelled secondary antibodies were developed using Clarity Western ECL substrate solutions (Bio-Rad) with a ChemiDoc XRS+ imaging system (Bio-Rad).

### Immunofluorescence microscopy and image analysis

Cells were seeded on glass coverslips, fixed with 4% formaldehyde (FA; 18814; Polysciences) for 12 min at room temperature, and permeabilized with 0.05% saponin (S7900; Sigma-Aldrich) in PBS. Fixed cells were then stained with primary antibodies at room temperature for 1 h in PBS/saponin, washed in PBS/saponin, stained with fluorescently labelled secondary antibody for 1 h, washed in PBS, and mounted with Mowiol containing 2 µg/ml Hoechst 33342 (H3570; Thermo Fisher Scientific).

For quantification, all images within one dataset were taken at fixed intensities below saturation, very high expressing cells have been omitted, and identical settings were applied for all treatments within one experiment. In general, 15 images were taken randomly from each condition.

The NIS-elements software was used for background correction (“rolling ball”) and automated image analyses. Identical analysis settings were applied for all treatments within one experiment. Fluorescent LC3B, p62 or WIPI2 dots were segmented by the software, and the number and sum fluorescence intensity of the dots were measured, as well as the co-occurrence of WIPI2 and DFCP1 dots. The total number of cells was quantified by automated detection of Hoechst nuclear stain by the software.

### Live Imaging of omegasome formation

Live-cell imaging was performed using either a DeltaVision OMX V4 microscope or a DeltaVision Elite deconvolution microscope. The DeltaVision OMX V4 was equipped with three PCO.edge sCMOS cameras, a solid-state light source, and laser-based autofocus, with environmental control provided by a heated stage and an objective heater (20/20 Technologies). The DeltaVision Elite was equipped with a 60×/1.42 NA oil-immersion objective and an sCMOS camera.

Movies were deconvolved using softWoRx software and processed in ImageJ/FIJI. Cells were imaged in EBSS, and images were taken every 2 sec over a period of 10-15 min. For imaging of SNAP-LC3B, cells were stained with SiR647-SNAP (New England Biolabs) according to the manufacturer’s protocol. Tracking of omegasomes and data processing was performed as previously described (Nahse et al., 2023).

### Statistics

The number of individual experiments and the number of cells or images analysed are indicated in the figure legends. The number of experiments was adapted to the expected effect size and the anticipated consistency between experiments. We tested our datasets for normal distribution by Kolmogorov-Smirnov, D’Agostino and Pearson, and Shapiro–Wilk normality tests, using GraphPad Prism Version 8. For parametric data, an unpaired two-sided t test was used to test two samples with equal variance, and a one-sample t test was used in the cases where the value of the control sample was set to 1. For more than two samples, we used ordinary one-way ANOVA with a suitable post hoc test. For nonparametric samples, Mann–Whitney test was used to test two samples and Kruskal–Wallis with Dunn’s post hoc test for more than two samples. All error bars denote mean values ±SD or SEM, as indicated in every figure legend (*, P < 0.05; **, P <0.01; ***, P < 0.001). No samples were excluded from the analysis.

## Acknowledgments

The X-ray diffraction was collected at the Swiss Light Source PX, Villigen, Switzerland and Diamond I04, UK beamlines. We thank the scientific staff at SLS PX and Diamond beamlines for their support. This work was supported by the DFG-funded Collaborative Research Center on Selective Autophagy (SFB1177) Project-ID 253130777 (I.D) and by the EUbOPEN Innovative Medicines Initiative 2 Joint Undertaking (JU) under grant agreement No. 875510 (M.M. and I.D.). We thank the Flow Cytometry Core Facility and the Advanced Light Microscopy Core Facility of Oslo University Hospital, and the Helsinki Institute of Life Science (HiLIFE) Light Microscopy Unit for technical assistance and access to instruments. We thank Deanna Wolfson and the Department of Physics and Technology, UiT The Arctic University of Norway, for access to and support with DeltaVision live-cell imaging.

V.N. was supported by a Mobility grant from the Research Council of Norway (grant number 301369) and from from the South-Eastern Norway Regional Health Authority (grant number 340630). K.O.S was supported by a Career grant from the South-Eastern Norway Regional Health Authority (grant number 2020038) and a Research Grant from the Research Council of Norway (grant number 315103). H.S. was supported by Project Grants from the South-Eastern Norway Regional Health Authority (project number 2018081) and the Norwegian Cancer Society (project number 182698), and by an Advanced Grant from the European Research Council (project number 788954). This work was partly supported by the Research Council of Norway through its Centres of Excellence funding scheme, project number 262652. H.S. and J.H.H. were supported jointly by a grant from the Peder Sather Foundation.

## Author Contributions

M.M., J.H.H., K.O.S., I.D., H.S. and V.N. designed the project and experiments. M.M., A.G. and J.P. designed constructs, performed mutagenesis and purified the ATPase constructs and mutants from bacteria while D.C. and S.M. purified the wild type ATPase domain from Sf9 insect cells. M.M., D.C. and S.M. performed protein crystallization and M.M. and S.M. solved the structures and concluded the modelling, refinement and PDB depositions. A.G. and M.M. performed the size exclusion runs and ATPase assay for ATPase wt and mutant proteins followed by analysis. Z.C. and K.T. purified the full-length DFCP1 from mammalian cells and performed GUV localisation assays. Y.Z. performed the ATPase assays for FL DFCP1 constructs. M-C.D.D., K.O.S. and V.N. generated cell lines, performed light microscopy and the omegasome formation assays in U2OS cells and carried out the analysis both for fixed and live cell imaging. M-C.D.D. and L.R.D.L.B performed analysis of fixed cell imaging. M.M., V.N. prepared the figures and wrote the manuscript with input from all the authors. J.H.H, K.O.S, I.D., H.S. and V.N. supervised the study, contributed to funding acquisition and research infrastructure, and edited the manuscript.

