## Supplemental figures for "Structural basis for the ATP-dependence of omegasome biogenesis by DFCP1"

### Supporting Information Text

### Figures

**Fig. S1. Crystal packing of DFCP1 ATPase domain in three crystal structures and different space groups**

(A) $DFCP1_{146\text{-}409}$ bound to ADP.Mg crystallized in P 1 21 1 spacegroup with 4 molecules (ChainA-D) in the asymmetric unit (AU). (B) $DFCP1_{146\text{-}409}$ bound to AppNHp crystallized in P 21 2 21 spacegroup with 3 molecules (Chain A, B, D) in the AU. (C) $DFCP1_{146\text{-}409}$ bound to ADP.Mg crystallized in C2 2 21 spacegroup with 2 molecules (Chain A, B) in the AU.

**Fig. S2. Multiple sequence aligment of DFCP1 ATPase domain (146-416) with the GTPase domain of Dynamin family proteins GBP1 (6-314), GBP2 (6-579), Atlastin1 (41-473), Atlastin2 (58-374), Atlastin3 (24-472) and IRGA6 (31-383) using Espript 3.0** (Gouet et al., 2003). The H195, R271, H323 and T369 positions of DFCP1 are highlighted in red boxes.

**Fig. S3. Multiple sequence aligment of DFCP1 ATPase domain (146-416) with the closely related ATPase domains of KIF5a (11-1020), p97 (159-691) and MYH2 (299-633) using Espript 3.0.** The H195, R271, H323 and T369 positions of DFCP1 are highlighted in red boxes.

**Fig. S4. Loss of the DFCP1 ATPase domain alters omegasome-phagophore dynamics**

(A) Quantification of WIPI2B-positive puncta per cell in DFCP1-knockout U2OS cells rescued with mNG-tagged DFCP1 WT or DFCP1 ΔATPase. Cells were starved in EBSS for 2 h, fixed, and stained for endogenous WIPI2B. Data are shown as mean ± SEM. (B) Quantification of WIPI2B-positive object area in DFCP1-knockout U2OS cells expressing mNG-DFCP1 WT or mNG-DFCP1 ΔATPase. Each point represents the median WIPI2B-positive object area per image; colours indicate independent experiments. (C) Representative images of cells treated as in A and B. Insets show magnified examples of DFCP1- and WIPI2B-positive omegasome-associated structures.

**Fig. S5. Omegasome dynamics are altered in arginine-finger and histidine-stack mutants.**

(A and B) Immunoblot analysis of DFCP1-knockout U2OS cells expressing mNG-tagged DFCP1 WT, DFCP1 ΔATPase, R271A, or H323A. Parental U2OS cells were included as a control. Actin was used as a loading control. (C) Representative time-lapse sequences of DFCP1-knockout U2OS cells co-expressing mNG-tagged DFCP1 WT, R271A, or H323A together with SNAP-LC3. Cells were starved in EBSS and imaged live by spinning-disk confocal microscopy to follow DFCP1-positive omegasomes and LC3-positive phagophores over time. Merged images are shown together with individual mNG-DFCP1 and SNAP-LC3 channels. Time is indicated in seconds. WT DFCP1-positive omegasomes expand into rings, separate from the SNAP-LC3-positive phagophore, and subsequently constrict. R271A-positive omegasomes undergo the same overall sequence but display delayed constriction. H323A-positive omegasomes show altered DFCP1–LC3 dynamics ('fading' phenotype).

**Fig. S6. Consurf (Ashkenazy et al., 2016) scores of DFCP1 amino acids mapped on the sequence and Alphafold3 predicted 3D structures.**

(A) Domain architecture of DFCP1 (1-777) molecule with colored domains. (B) The primary sequence of DFCP1 colored according to the conservation score (dark maroon: most conserved; dark teal: most variable). Residues 11-145 correspond to N-terminal domain, ATPase domain spans residues 146-416, ER-targeting domain connects ATPase domain with FYVE domain I (598-659) and continues to terminate at FYVE domain II (715-777). (C) Alphafold3 (Abramson et al., 2024) prediction of DFCP1 ATPase domain dimer with ATP and $Mg^{2+}$ (in spheres). (D) Superimposition of Alphafold3 prediction with AppNHp bound DFCP1 ATPase domain dimer in the crystal structure. The RMSD of the Alphafold3 prediction and crystal structure is 0.677 Å. (E) Surface representation of Alphafold3 predicted monomer colored based on the Consurf scores bound to ATP with two views.

**Fig. S7. Summary model of altered omegasome morphology in DFCP1 ATPase-domain mutants.**

Schematic summary of omegasome-phagophore dynamics observed by live-cell imaging in cells expressing DFCP1 WT, ΔATPase, R271A, or H323A. In WT cells, DFCP1-positive omegasomes expand into rings around LC3-positive phagophores, separate from the LC3-positive structure, and subsequently constrict. ΔATPase-positive omegasomes frequently fail to separate from LC3-positive structures and instead remain associated with the phagophore, often showing transient tubulation and a 'fading' phenotype. R271A-positive omegasomes undergo the canonical sequence of expansion, separation, and constriction, but constriction is delayed and the maximum omegasome diameter is reduced. H323A-positive omegasomes resemble the ΔATPase phenotype, with impaired DFCP1–LC3 separation, frequent fading, and tubulation. Green indicates DFCP1-positive omegasomes; magenta indicates LC3-positive phagophores.

**A**

**DFCP1 ATPase domain (146-409) + ADP.Mg: P 1 21 1 (4 mol/ ASU)**

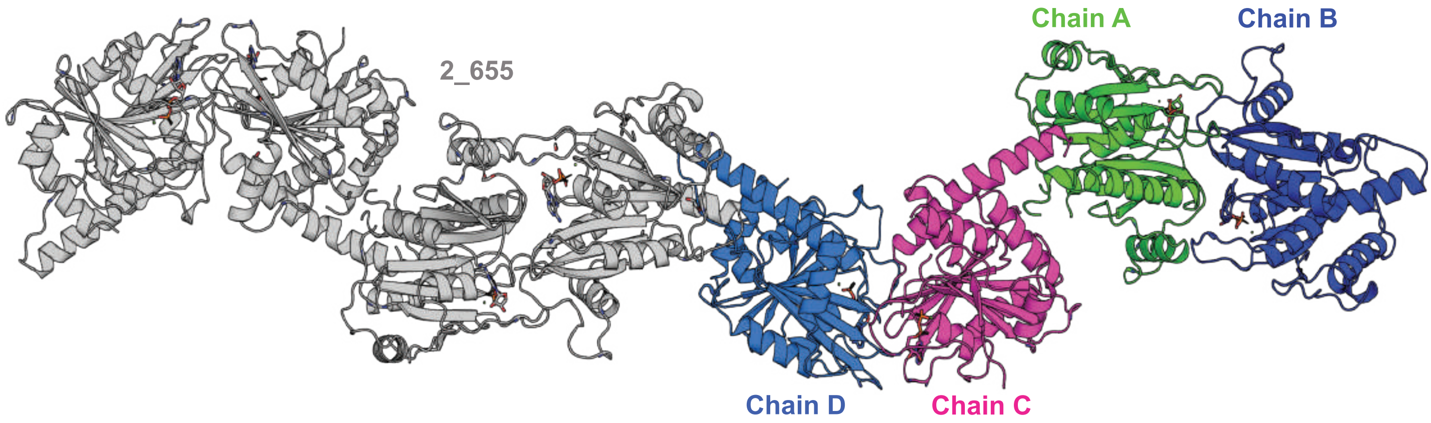


**B**

**DFCP1 ATPase domain (146-409) + AppNHp.Mg: P 21 2 21 (3 mol/ ASU)**

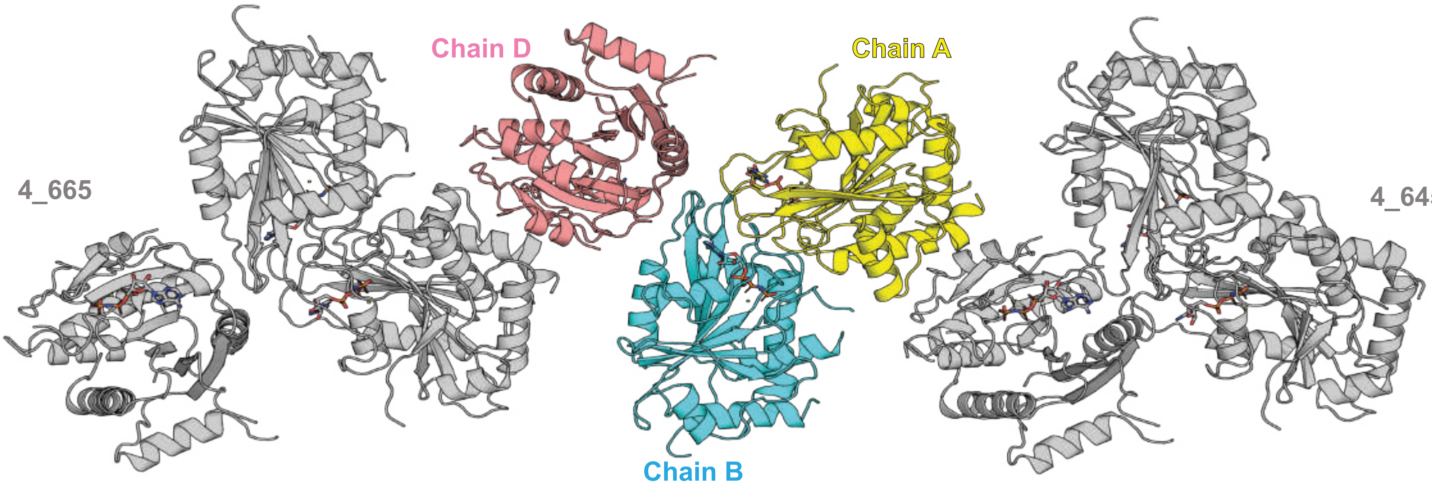


**C**

**DFCP1 ATPase domain (146-409) + ADP.Mg: C2 2 21 (2 mol/ ASU)**

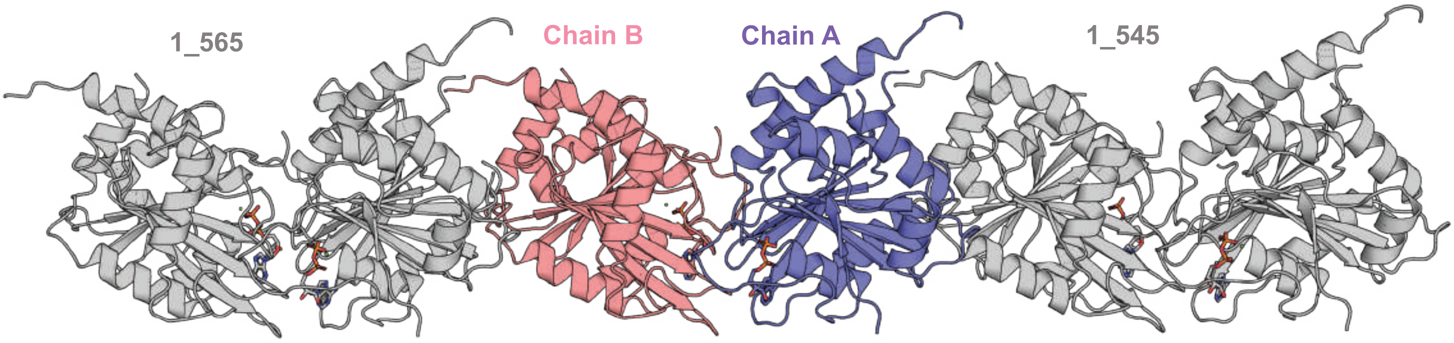


***SI Appendix*, Fig. S1**

***SI Appendix*, Fig. S2**

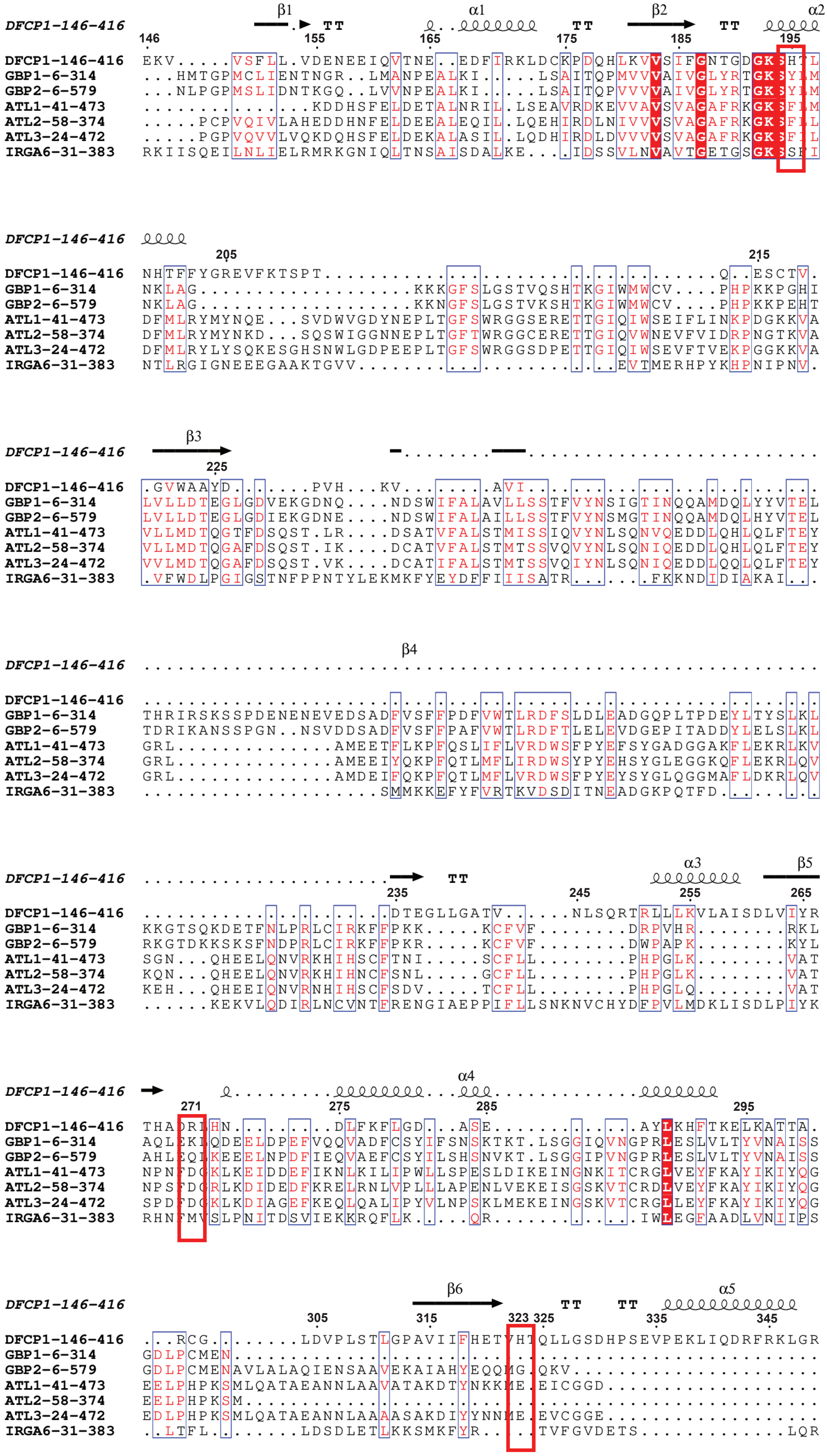

***SI Appendix,* Fig. S2**

```
DFCP1-146-416    TT     β7                α6                             
                   355       365 369    375       385       395
DFCP1-146-416    FPEAFSSIHYKGTRTYNPPTDFSGL........RRALEQLLENN....TTRSPRHPGVIF
GBP1-6-314       ............................................................
GBP2-6-579       ...............QLPTETLQELLDLHRDSEREAIEVFMKN...SFKDVDQMFQRKLG
ATL1-41-473      .............KPFLAPNDLQ...TKHLQLKEESVKLFRGVKKMGGEEFSRRYLQQLE
ATL2-58-374      ............................................................
ATL3-24-472      .............KPYLSPDILE...EKHCEFKQLALDHFKKTKKMGGKDFSFRYQQELE
IRGA6-31-383     LAR.................DWEIE........VDQVEAMIKSP....AVFKPTDEETIQ

                 α7
DFCP1-146-416
                          405       415
DFCP1-146-416    KALKALSDRFS....GEIPDDQM.....................................
GBP1-6-314       ............................................................
GBP2-6-579       AQLEARRDDFCKQNSKASSDCCMALLQDIFGPLEEDVKQGTFSKPGGYRLFTQKLQELKN
ATL1-41-473      SEIDELYIQYIKHNDSKNIFHAARTPATLFVVI.........................FI
ATL2-58-374      ............................................................
ATL3-24-472      EEIKELYENFCKHNGSKNVFSTFRTPAVLFTGI.........................VA
IRGA6-31-383     ERLSRYIQEFCLANGYLLPKNSF.....................................

DFCP1-146-416

DFCP1-146-416    ............................................................
GBP1-6-314       ............................................................
GBP2-6-579       KYYQVPRKGIQAKEVLKKYLESKEDVADALLQTDQSLSEKEKAIEVERIKAESAEAAKKM
ATL1-41-473      TYVIAGVTGFIGLDII............................................
ATL2-58-374      ............................................................
ATL3-24-472      LYIASGLTGFIGLEVVAQ..........................................
IRGA6-31-383     ............................................................

DFCP1-146-416

DFCP1-146-416    ............................................................
GBP1-6-314       ............................................................
GBP2-6-579       LEEIQKKNEEMMEQKEKSYQEHVKQLTEKMERDRAQLMAEQEKTLALKLQEQERLLKEGF
ATL1-41-473      ............................................................
ATL2-58-374      ............................................................
ATL3-24-472      ............................................................
IRGA6-31-383     ............................................................

DFCP1-146-416

DFCP1-146-416    ................
GBP1-6-314       ................
GBP2-6-579       ENESKRLQKDIWDIQM
ATL1-41-473      ................
ATL2-58-374      ................
ATL3-24-472      ................
IRGA6-31-383     ................
```

**SI Appendix, Fig. S3**

```
DFCP1-146-416   β1 TT
                146 155
DFCP1-146-416   ........................................EKVVSFLLVDENEEIQ....
KIF5A-11-1020   .......................KVLCRFRPLNQAEILRGDKFIPIFQG...DDSV....
p97-159-691     MRAVEFKVVETDPSPYCIVAPDTVIHCEGEPIKRE..............DEEESLNEVGY
MYH2-299-633    .ELIEMLLITTNPYDYPFVSQGEISVASI..DDQEELMATDSAIDILGFTNEEKVS..IY

DFCP1-146-416   α1 TT β2
                165 175 185
DFCP1-146-416   .VTNEE.........DFIRK................LD..CKPDQ.......HLKVVSIF
KIF5A-11-1020   .VIGGK.........PYVFDRVFPPNTTQEQ...VYHA..CAMQIVKDVLAGYNGTIFAY
p97-159-691     DDIGGC.........RKQLAQIKE..MVELP...LRH.....PALFKAIGVKPPRGILLY
MYH2-299-633    KLTGAVMHYGNLKFKQKQREEQAEPDGTEVADKAAYLQSLNSADLLKAL..CYPRV..KV

DFCP1-146-416   TT α2
                195 205
DFCP1-146-416   GNTGDGKSHTLNHTF..........FYG..........REVFK.................
KIF5A-11-1020   GQTSSGKTHTMEGKL..........HDPQLMGIIPRIARDIFNHIYSMDENLEFHIKVSY
p97-159-691     GPPGTGKTLIARAVANETGAFFFLINGPEIMSKLAGESESNLRKAFEEA......EKNAP
MYH2-299-633    GNEYVTKGQTVEQVSNAVGALAKAVYEKMFLWMVARINQ...........QLDTKQPRQY

DFCP1-146-416   β3
                215 225 235
DFCP1-146-416   ..........................TSPTQESCTVGVWAAYDPVHKVAVID........
KIF5A-11-1020   FEIYLDKIRDLLDVTKTNLSVHEDKNRVPFVKGCTERFVSS..PEEILDVIDEGKSNRHV
p97-159-691     AIIFIDELDAIAPKRE...K..........THGEVERRI....VSQLLTLMDGLKQRAHV
MYH2-299-633    FIGVLDI....AGFEI.............FDFNSLEQLCINFTNEKL.............

DFCP1-146-416   β4 TT
                245
DFCP1-146-416   ...............................................TEGLLGATVNLSQ
KIF5A-11-1020   AVTNMNEHSSRSHSIFLINIKQENMETEQKLSGKLYLVDLAGSEKVSKTGAEGAVLDEAK
p97-159-691     IVMAA..............TN....................................RPN
MYH2-299-633    .......QQFFNHHMFVLEQ.......................EEYKKEGIE........

DFCP1-146-416   α3 β5
                255 265
DFCP1-146-416   R....TRLLLKV...LA................ISDLVIYR...................
KIF5A-11-1020   NINKSLSALGNVISALAEG..............TKSYVPYRDSKMTRILQDSLGGNCRTT
p97-159-691     SIDPALRRFGRFDREVDIGIPDATGRLEILQIHTKNMKLADDVDLEQVANE.THGHVGAD
MYH2-299-633    ............................................................

DFCP1-146-416
                271 275
DFCP1-146-416   ......................THADRLHNDLFKFL..GDAS..................
KIF5A-11-1020   MFICCSPSSYNDAETKSTLMFGQRAKTIKNTASVNL..ELTAEQWKKKYEKEKEKTKAQK
p97-159-691     LAALCSEAALQAIRKKMDLI.DLEDETIDAEVMNSLAVTMDDFRWALSQSN....PSALR
MYH2-299-633    ............................................................

DFCP1-146-416   ............................................................

DFCP1-146-416   ............................................................
KIF5A-11-1020   ETIAKLEAELSRWRNGENVPETERLAGEEAALGAELCEETPVNDNSSIVVRIAPEERQKY
p97-159-691     ETV..VEVPQVTWEDIGGL.........................................
MYH2-299-633    ............W...............................................

DFCP1-146-416   ............................................................

DFCP1-146-416   ............................................................
KIF5A-11-1020   EEEIRRLYKQLDDKDDEINQQSQLIEKLKQQMLDQEELLVSTRGDNEKVQRELSHLQSEN
p97-159-691     ...............................................EDVKRELQE....
MYH2-299-633    ............................................................

DFCP1-146-416   ............................................................

DFCP1-146-416   ............................................................
KIF5A-11-1020   DAAKDEVKEVLQALEELAVNYDQKSQEVEEKSQQNQLLVDELSQKVATMLSLESELQRLQ
p97-159-691     ............LVQYPVEHP.......................................
MYH2-299-633    ............................................................
```

### *SI Appendix*, Fig. S3

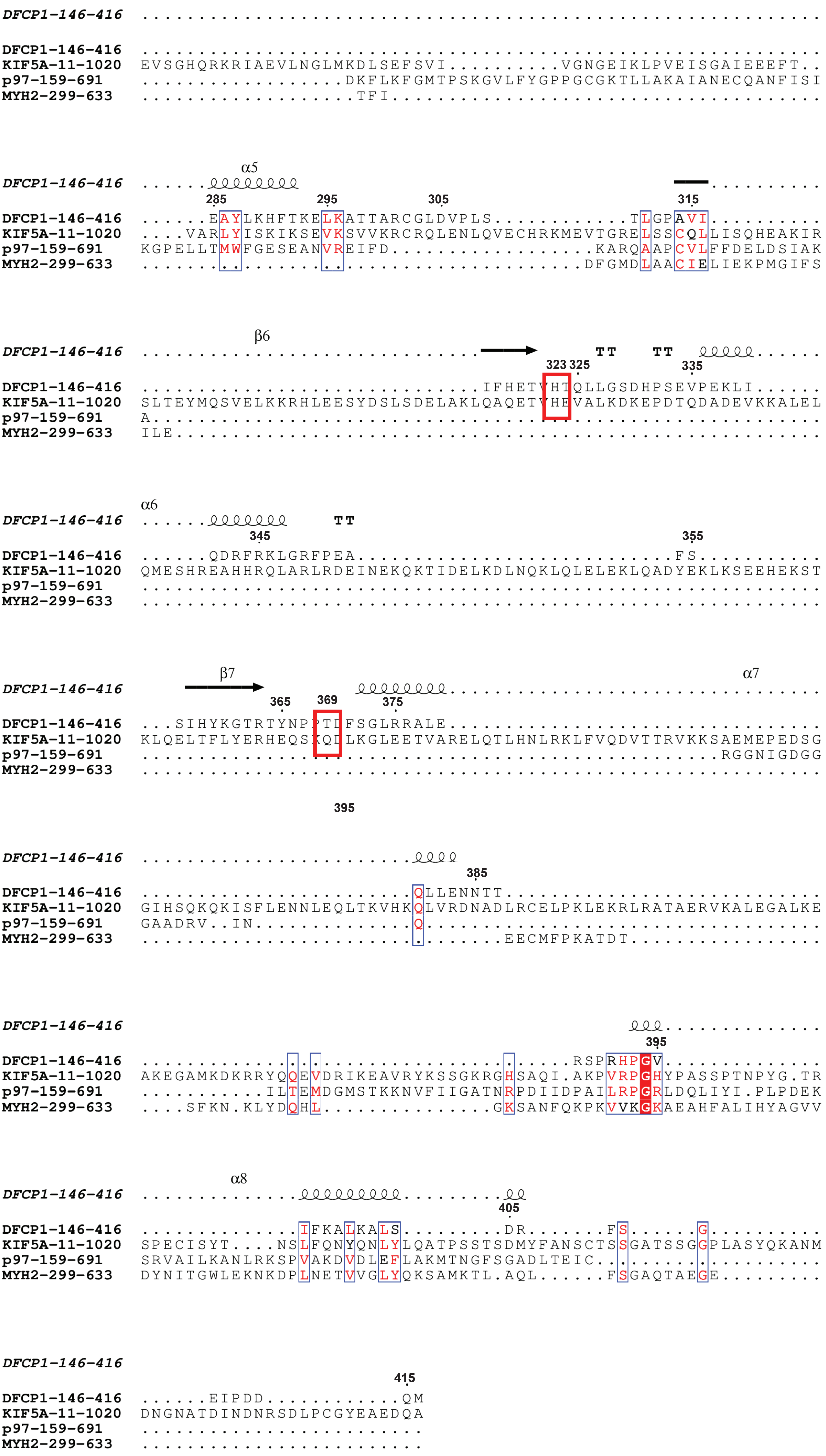

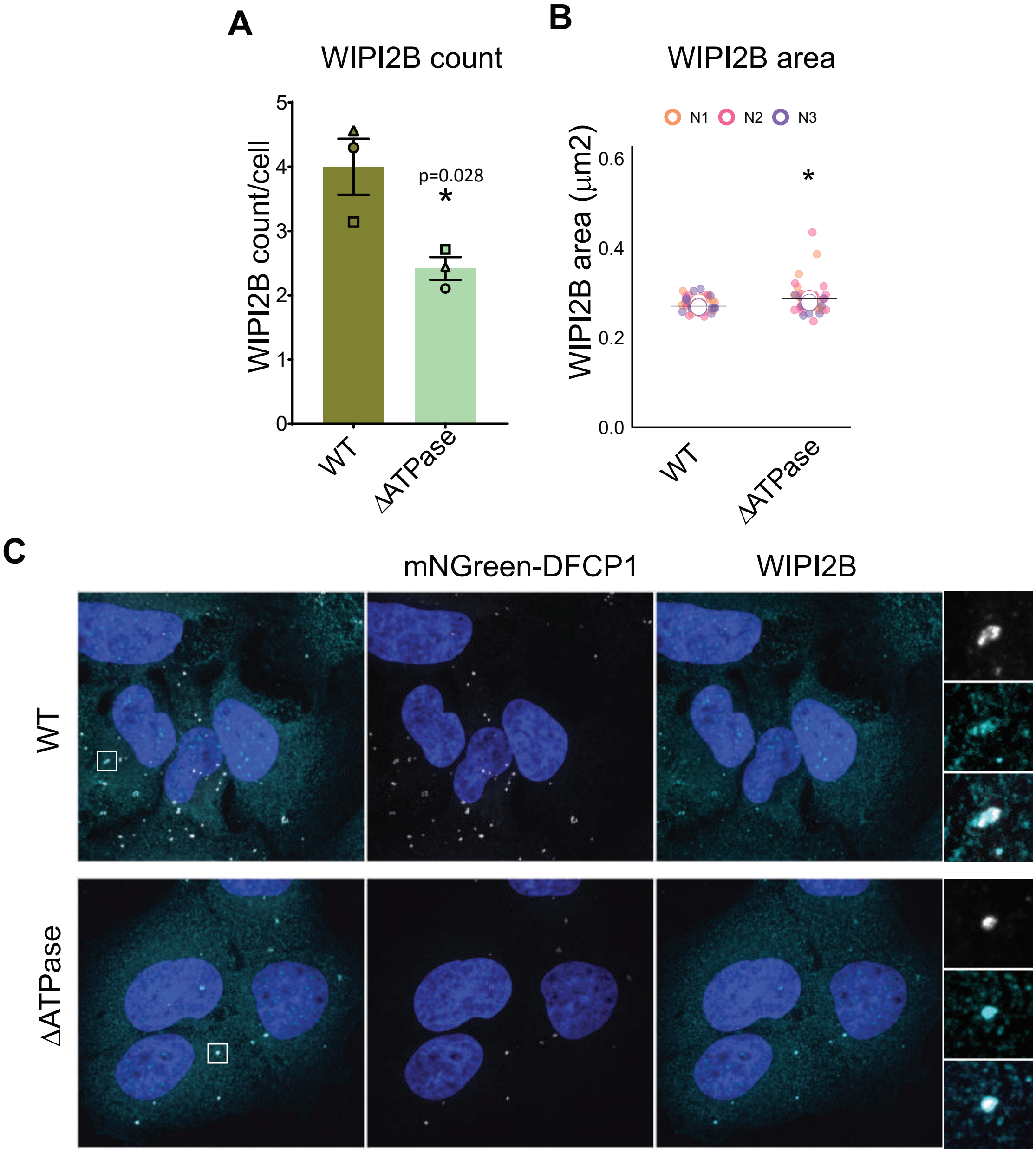


***SI Appendix*, Fig. S4**

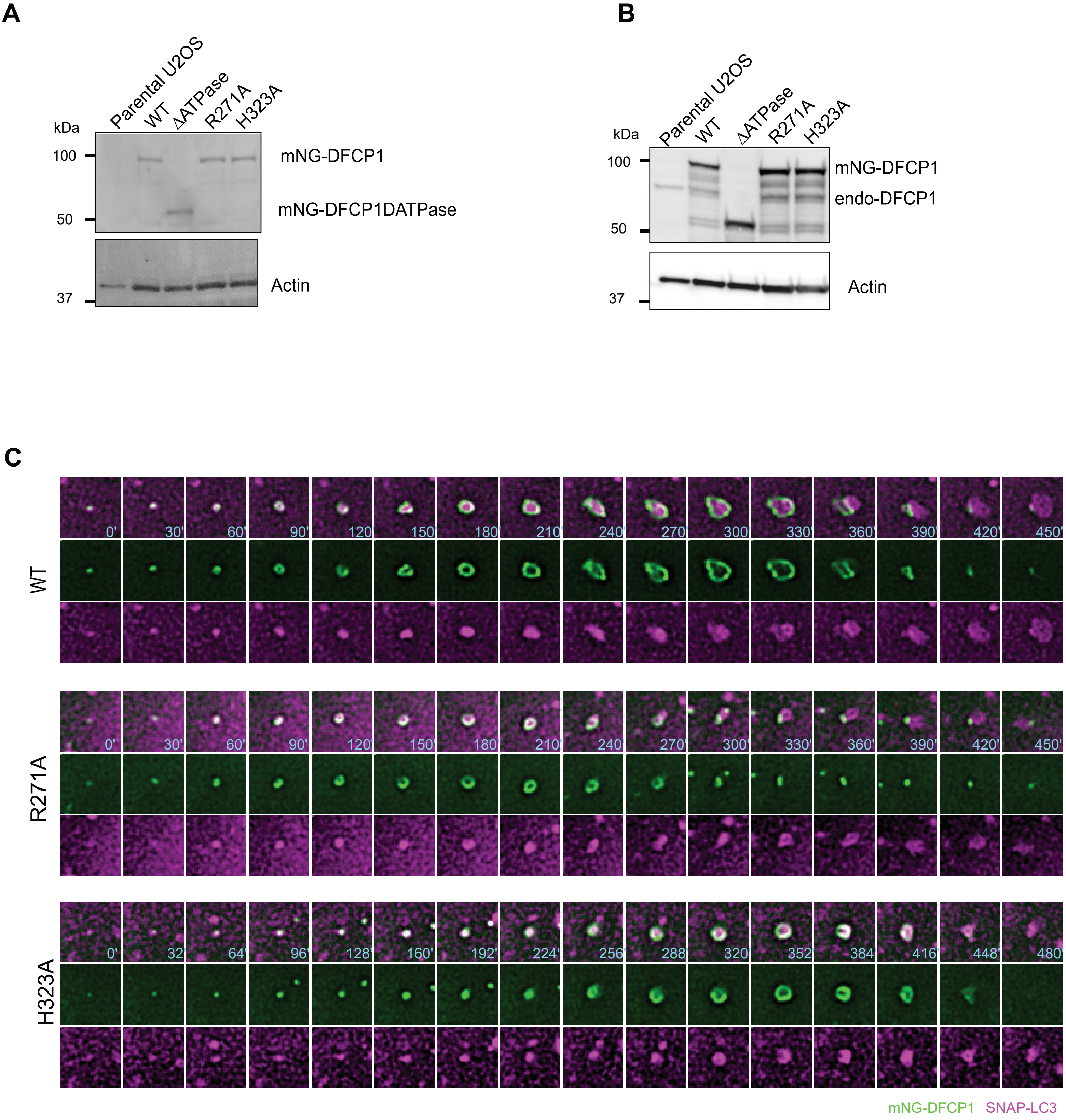


***SI Appendix*, Fig. S5**

### SI Appendix, Fig. S6

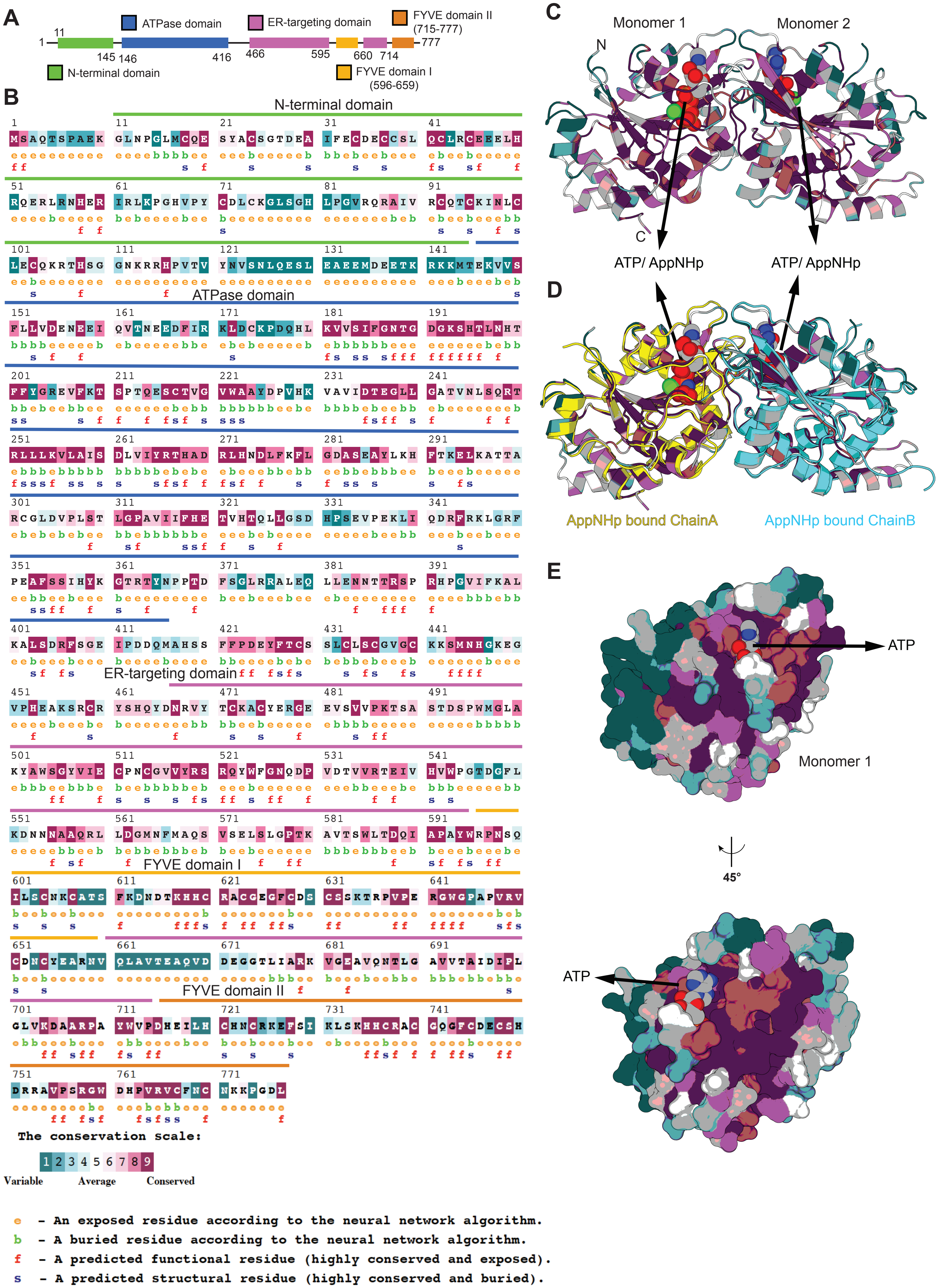

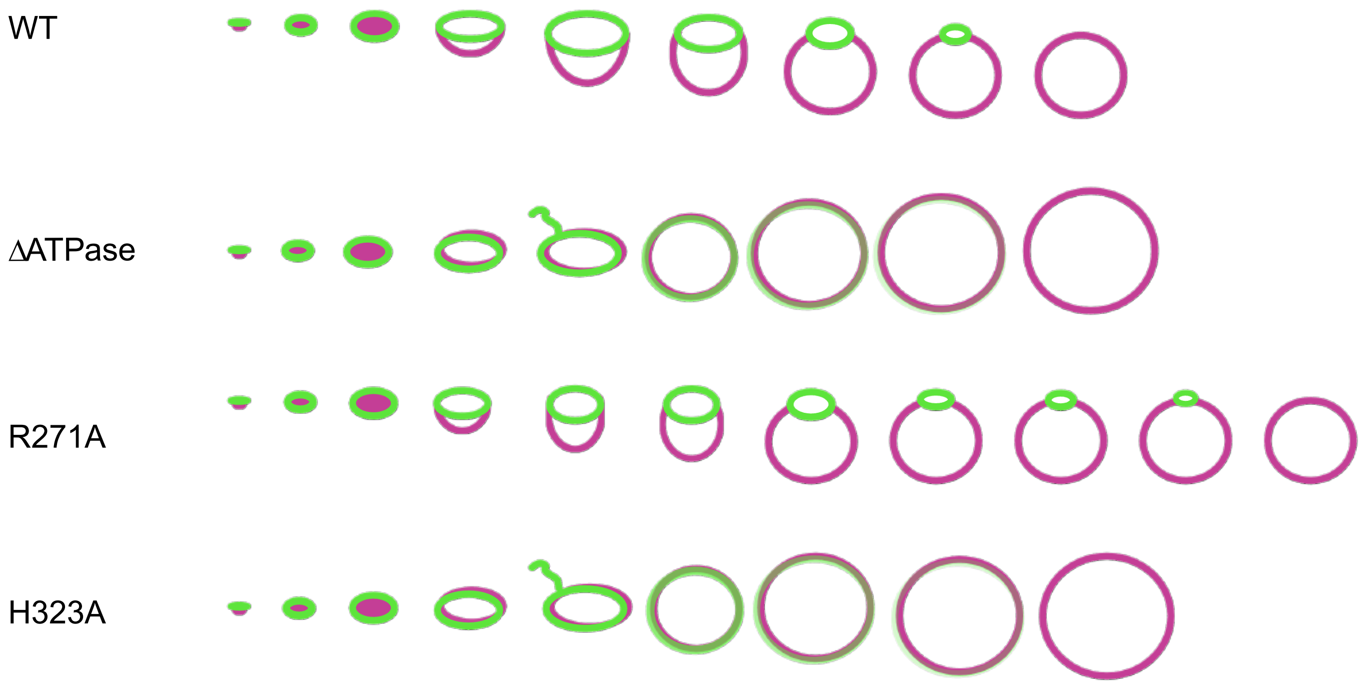


***SI Appendix*, Fig. S7**
